# *VGLL2-NCOA2* leverages developmental programs for pediatric sarcomagenesis

**DOI:** 10.1101/2021.09.15.460544

**Authors:** Sarah Watson, Collette LaVigne, Lin Xu, Whitney Murchison, Didier Surdez, Dinesh Rakheja, Franck Tirode, Olivier Delattre, James Amatruda, Genevieve Kendall

## Abstract

Clinical sequencing efforts are uncovering numerous fusion genes in childhood solid tumors, yet few methods exist to delineate fusion-oncogenes from structural changes of unknown significance. One such novel fusion gene is *VGLL2-NCOA2*, which was described by us and others in patients with congenital sarcomas but has lacked functional validation. To determine if this fusion is an oncogene, and how it is driving disease, we developed a vertebrate zebrafish model and mouse allograft model of human *VGLL2-NCOA2* driven sarcomagenesis. We found that *VGLL2-NCOA2* is indeed an oncogene and is sufficient to generate mesenchymal tumors that recapitulate the human disease at the histological and transcriptional level. Zebrafish *VGLL2-NCOA2* tumors display features of arrested skeletal muscle development, and a subset transcriptionally cluster with somitogenesis in developing embryos. By comparing tumor and embryonic gene expression signatures, we identified developmentally regulated targets that *VGLL2-NCOA2* potentially leverages for tumorigenesis. These targets highlight the core biology of the disease and could represent therapeutic opportunities. Specifically, a RAS family GTPase, *arf6/ARF6*, involved in actin remodeling and rapid cycling of endocytic vesicles at the plasma membrane, is highly expressed during zebrafish somitogenesis and in *VGLL2-NCOA2* tumors. In zebrafish tumors, arf6 protein is highly expressed and is absent from mature skeletal muscle. In *VGLL2-NCOA2* mouse allograft models and patient tumors, *ARF6* mRNA is overexpressed as compared to skeletal muscle or normal controls. More broadly, *ARF6* is overexpressed in adult and pediatric sarcoma subtypes as compared to mature skeletal muscle. Overall, our cross-species comparative oncology approach provides evidence that *VGLL2-NCOA2* is an oncogene which leverages developmental programs for tumorigenesis, and that one of these programs, the reactivation or persistence of arf6/ARF6, could represent a therapeutic opportunity.

## Introduction

With the broader implementation of clinical sequencing efforts, there has been a significant increase in the number of gene fusions that have been identified in patient tumors. Particularly in sarcomas, there is a predilection for fusion-oncogenes with a predicted 20-49% of sarcomas containing a gene fusion event ^1–4^. In pediatric sarcomas, there are typically few other somatic mutations, thus highlighting fusions as the defining oncogenic drivers in these diseases ^5–7^. However, developing cell culture or animal models of these newly identified fusions is challenging, hindering the verification of their role in tumorigenesis and the identification of potential targeted therapeutics. A pressing need exists for tractable genetic models to understand the shared and divergent biology between genetic drivers, and ultimately provide a platform to identify mechanism based therapies.

One example of a newly discovered fusion gene is *VGLL2-NCOA2*, which was identified from a cohort of congenital pediatric sarcomas that did not contain the classical pediatric sarcoma gene fusions ^4,8^. Both VGLL2 and NCOA2 are transcriptional co-activators, but their function in this gene fusion has not been studied. VGLL2 (previously named VITO-1) plays a role in muscle development and targeting TEAD proteins to their appropriate contexts during the differentiation process ^9–11^. Additionally, in zebrafish models, *vgll2a* is required for neural crest cell survival and craniofacial development ^12^. *NCOA2* is a common 3’ fusion partner in non-spindle cell sarcoma contexts; examples including *HEY1-NCOA2* in mesenchymal chondrosarcoma, *MEIS1-NCOA2* in genitourinary and gynecologic tract spindle cell sarcomas and intraosseous rhabdomyosarcomas, and *PAX3-NCOA2* in alveolar rhabdomyosarcoma ^13–17^. Suggestively, *TEAD1-NCOA2* fusions have also been identified in infantile rhabdomyosarcoma, indicating converging developmental processes for this subtype ^18,19^.

The *VGLL2-NCOA2* fusion has been described in multiple patients by multiple groups. There are two forms of the gene fusion, including *VGLL2* exons 1-2 and either *NCOA2* exons 13-23 or 14-23. The translocation involves two genomic events including an inversion of *VGLL2*, followed by a break and chromosomal translocation of chromosomes 6 and 8. These fusions are the defining genetic features of the disease; however, pathologically and transcriptionally, the tumors are heterogeneous ^4,8,20^. The biological explanation for this heterogeneity remains unclear.

Clinically, *VGLL2-NCOA2* tumors typically present between 0-1 year of age and have a spindle and sclerosing cellular morphology ^4,8,18,20–22^. Fusion-driven congenital rhabdomyosarcomas generally have a favorable prognosis with successful surgery and chemotherapy ^8^. However, non-resectable tumors are challenging to treat, and chemotherapy exposure carries a significant risk of short and long-term adverse effects. Moreover, clinical outcomes of *VGLL2-NCOA2* rearranged tumors are variable. Recently, a case study with four *VGLL2* rearranged tumors described three patients that had multi-metastatic spread, and two that died from disease. This study suggests that complete surgical resection is critical (however not always attainable) for long-term benefit, and that *VGLL2* fusions are capable of aggressive disease ^22^. Given that these tumors occur at a developmentally sensitive age, effective targeted therapies are critically needed to improve disease outcomes and ameliorate off-target effects from toxic and generalized therapies. Therefore, we set out to test the tumorigenic capacity of this novel fusion gene, define its tumorigenic program, and identify rational therapeutic targets.

Previously, we described a mosaic strategy using vertebrate transgenic zebrafish models that addresses the need to readily generate animal models that recapitulate the genetics and presentation of the human disease. This strategy has been successful for *PAX3-FOXO1* driven rhabdomyosarcoma and *CIC-DUX4* mediated Ewing-like sarcoma, suggesting that zebrafish are a powerful tool for personalized medicine approaches and for the study of rare cancers and disease ^23,24^. Here, we implement a functional genomics approach to generate a human *VGLL2-NCOA2* fusion-oncogene driven zebrafish and mouse allograft models of tumorigenesis. We find that the fusion is indeed sufficient for aggressive tumorigenesis in the zebrafish and mouse systems, resulting in tumors that recapitulate features of the human disease. Zebrafish *VGLL2-NCOA2* tumors express markers indicative of arrested muscle development and poor differentiation. Sequencing of zebrafish and mouse allograft tumors identified ARF6, a small GTPase, as an overexpressed and potentially druggable target. ARF6 is similarly expressed not only in human *VGLL2-NCOA2* tumors but also across a panel of pediatric sarcomas and other cancers, whereas there is no expression in their normal tissue counterparts. This study suggests that fusion-driven congenital rhabdomyosarcomas co-opt developmental programs to induce their effects and that zebrafish are high fidelity models of these systems. Further, these models have the potential to identify novel tumor-specific candidate therapeutic targets that may be relevant across a range of pediatric cancers.

## Results

### VGLL2-NCOA2 is an oncogene and is tumorigenic in zebrafish systems

Previously, we and others have shown that *VGLL2-NCOA2* is a novel fusion gene identified in a subset of congenital rhabdomyosarcomas ^4,8^. Five patients under the age of five presented with soft tissue tumors that were originally diagnosed as embryonal rhabdomyosarcomas. Clinical sequencing of patient tumors identified the presence of a *VGLL2-NCOA2* fusion. Supplemental Table 1 indicates the clinical presentation of each case that were identified in Watson et al 2018 ^4^. The fusion in all tumors contains an intronic breakpoint in *VGLL2* that results in the incorporation of the first two exons of *VGLL2*. However, there were two intronic breakpoints seen in *NCOA2*; four tumors had a breakpoint that resulted in inclusion of exons 14 through 23 of *NCOA2*, whereas one tumor showed inclusion of exons 13-23 (Supplemental Table 1; Supplemental Figure 1) ^4^.

The coding sequence of the human fusion was cloned out of a primary patient tumor that harbored the *NCOA2* exon 14 breakpoint (Supplemental Figure 1A). Gateway cloning was used to generate a CMV-GFP2A-*VGLL2NCOA2* construct for expression of the human fusion gene in zebrafish systems. The cassette is flanked on both sides by Tol2 sites allowing for genomic integration when injected with Tol2 mRNA (Figure 1A) ^25^. The viral 2A sequence allows for GFP and *VGLL2-NCOA2* to be transcribed on the same mRNA, yet function as independent proteins. We injected this expression construct and Tol2 mRNA into one-cell stage wildtype zebrafish embryos and observed robust GFP expression at 24 hours post fertilization in >95% of embryos (data not shown). Zebrafish were monitored for tumor development by observing gross morphology and GFP expression, which served as a proxy for fusion-oncogene expression. Zebrafish presented with tumors as early as one month, with 20% of zebrafish developing tumors by 50 days, and ~30% by six months. In control cohorts injected with CMV-GFP-pA, tumors were never observed (Figure 1B). The sufficiency of the *VGLL2-NCOA2* fusion to generate tumors in zebrafish confirms its function as an oncogene.

**Figure 1.**
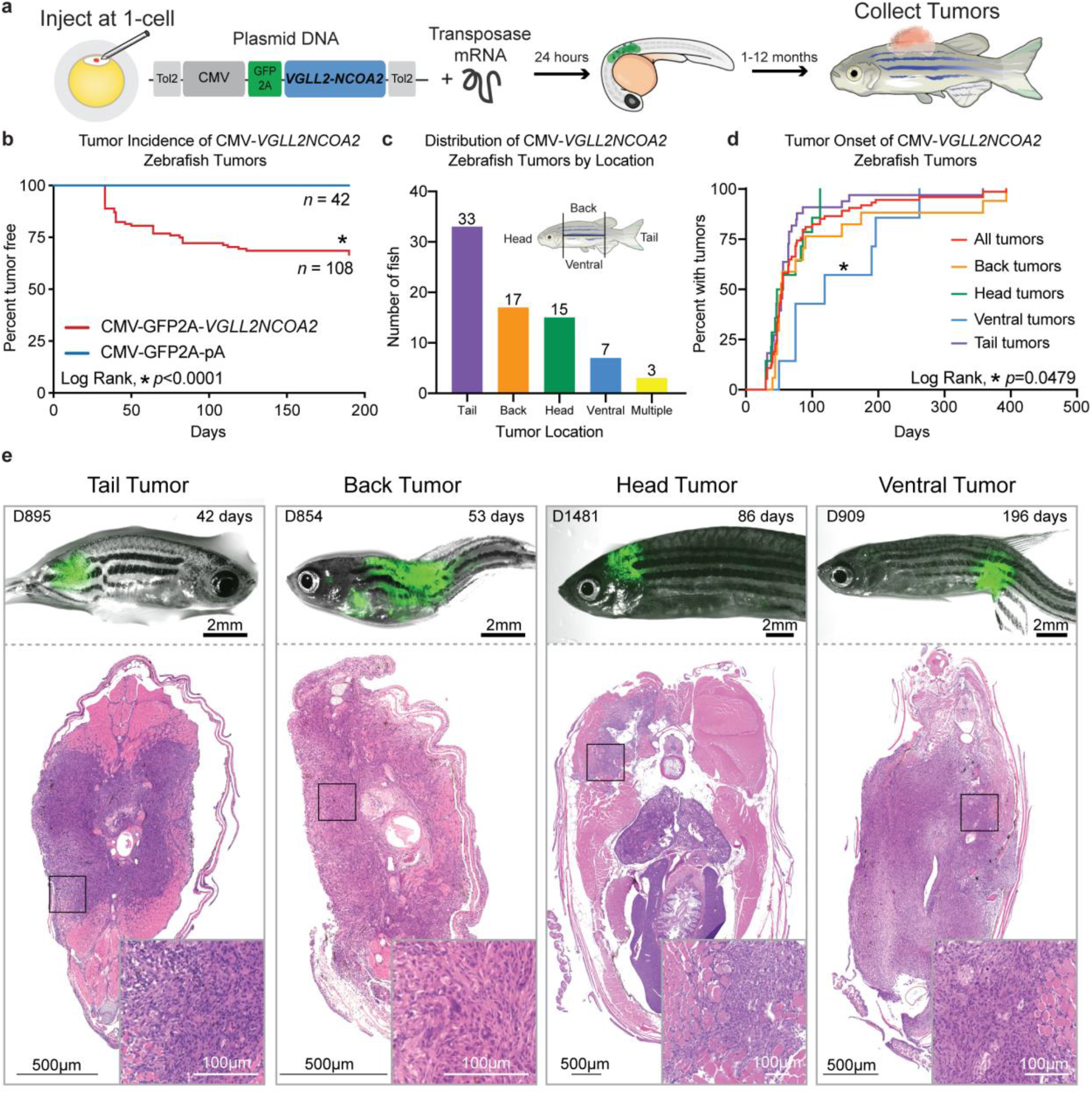
A new zebrafish model of human *VGLL2-NCOA2* driven tumorigenesis. A) Zebrafish were injected at the 1-cell stage with plasmid DNA carrying the human fusion gene *VGLL2-NCOA2* driven by the CMV promoter and tagged with GFP2A and Tol2 transposase mRNA to stably integrate the construct. The GFP+ embryos were tracked from 1-12 months for tumor formation. B) Tumor incidence curve of one independent experiment showing *n*=108 adult fish injected with CMV-GFP2A-*VGLL2NOCA2* compared to n=42 adult sibling fish injected with CMV-GFP2A-pA and Log Rank (Mantel-Cox) test performed with *p*<0.0001. All fish that survived past 30 days of age are included. This experiment was repeated in three independent cohorts, with similar tumor incidence curves (data not shown). C) Distribution of CMV-*VGLL2NCOA2* tumors classified by location as depicted on the inset fish schematic. D) Tumor onset curve for fish with a CMV-*VGLL2NCOA2* driven tumor, stratified by location on the fish as defined in C. Log Rank (Mantel-Cox) analysis showed significant difference between the ventral tumors and all other tumors (*p*=0.0479). E) Representative CMV-*VGLL2NCOA2* injected fish with tumors classified by location. For each fish, brightfield and GFP fluorescent images are overlaid and shown above the hematoxylin and eosin stain of a transverse section through the GFP+ area. The top left inset is the unique fish ID, and the top right is the age of the fish when the tumor was resected.

We classified all generated tumors by location on the fish, binning the fish as depicted in Figure 1C. We observed that tumors arising on the tail were the most common (44%; n=33/75), back and head tumors occurred in equal numbers (23%; n=17/75 and 20%; n=15/75 respectively), and ventral tumors were the least common (9%; n=7/75) (Figure 1C). The tumor onset was similar for each location with the exception of the ventral tumors which arose later compared to the other tumor locations (Log-Rank, p<0.05) (Figure 1D). Interestingly, three zebrafish presented with more than one tumor around 47 days of age (Figure 1C, Figure 3C and Supplemental Table 2).

Shown in Figure 1E is the gross morphology and GFP fluorescence of four representative zebrafish tumors and their corresponding hematoxylin and eosin (H&E) stains of transverse sections through the GFP+ region. All four of these tumors are mesenchymal tumors diagnosed as sarcomas, with infiltrating cells, and a variety of cells including spindle cells, round cells and pleomorphic cells, though in differing proportions. All tumors are mitotically active. The tumors contain dispersed nuclear chromatin and poorly defined cell membranes; prominent nucleoli are seen in the head tumor. Fascicular arrangements of cells are observed in the tail and back tumors, and there is a notable lack of background stroma. In the ventral tumor, there is a higher proportion of stroma and more pleomorphic and spindle cells. Collagen is present in variable amounts in the ventral tumor. In the head tumor, the predominant cell types are pleomorphic and round cells with accompanying necrosis. Altogether, these tumors are consistent with the histology of human sarcomas.

We also generated constructs in which GFP-tagged *VGLL2-NCOA2* was driven by the following zebrafish promoters: MCS-betaglobin-Splice Acceptor, beta-actin, ubiquitin, and unc503. Following zebrafish microinjection, we observed GFP expression at 24 hours post-fertilization, indicating successful genomic integration of the constructs (Supplemental Figure 2A). With MCS-betaglobin-SA-GFP2A-*VGLL2NCOA2*, we observed a total of three fish out of 42 that presented with tumors, two diagnosed as sarcomas and one as neuroblastoma/retinoblastoma (Supplemental Figure 2B). The other promoters failed to produce any tumors, though some GFP+ adult fish were observed (Supplemental Figure 2C). Given the low incidence, we focused our efforts on the CMV-driven model which recapitulated the human disease with the highest fidelity. All zebrafish tumors in the remainder of the study are generated from the CMV-GFP2A-*VGLL2NCOA2* construct.

### VGLL2-NCOA2 zebrafish tumors recapitulate the human disease

To validate that zebrafish tumors expressed the VGLL2-NCOA2 fusion at the protein level we performed immunohistochemistry on serial transverse sections of zebrafish tumors using a human NCOA2 antibody (Figure 2A). The tumor cells identified with H&E staining were also positive for NCOA2. Additionally, zebrafish mature skeletal muscle was negative for NCOA2, indicating that VGLL2-NCOA2 is specifically expressed in the GFP positive tumors. We confirmed by RT-PCR that *VGLL2-NCOA2* tumors expressed the fusion at the mRNA level (Figure 2B). Further, zebrafish *VGLL2-NCOA2* tumors expressed the muscle markers *myod1, myog*, and *desma*, which are the zebrafish orthologs of *MYOD, MYOG* and *DESMIN*. These muscle markers are used to clinically characterize the human disease (Figure 2B; Supplemental Table 1) ^4,8^.

**Figure 2.**
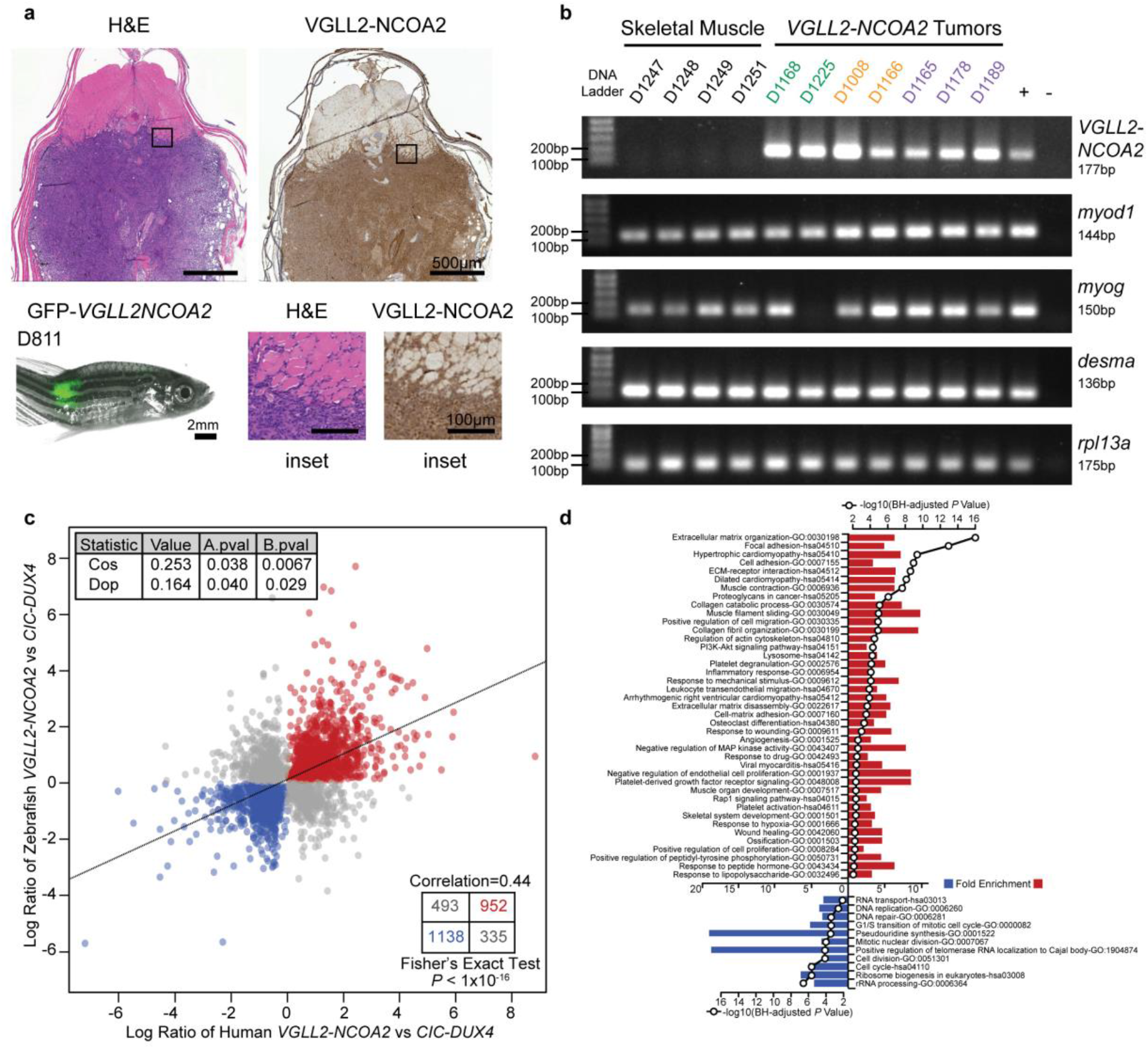
Zebrafish *VGLL2-NCOA2* tumors recapitulate the human disease. A) Zebrafish CMV-*VGLL2NCOA2* tumor shown with GFP expression and gross morphology. Transverse sections through the GFP+ region were stained with hematoxylin and eosin, and the subsequent step section stained with an anti-NCOA2 antibody. All n=9 zebrafish tumors stained were positive. B) RT-PCR on RNA isolated from zebrafish *VGLL2-NCOA2* tumors and adult zebrafish skeletal muscle showing presence or absence of the oncogene and muscle markers: *myod1, myog, desma* and *rpl13a* control. Tumors are colored based on location on the fish: back tumors in orange, head tumors in green and tail tumors in purple. C) RNA sequencing data of *VGLL2-NCOA2 (n=11)* and *CIC-DUX4* (n=12; from Watson et al 2019 ^24^) driven zebrafish tumors were compared to their human counterpart (*VGLL2-NCOA2* n=8 and *CIC-DUX4* n=11; from Watson et al 2018 ^4^) in an Agreement of Differential Expression analysis. Only the 2918 genes that were differentially expressed between the two tumor types in either sets of data are plotted. The different statistics of the analysis are shown as well as the Pearson correlation coefficient and the exact Fisher test P-value of the shown contingency table. D) Gene ontology terms associated with the n= 952 genes up- or n=1138 down-regulated in *VGLL2-NCOA2* tumors. Fold enrichment (barplot) and adjusted BH P-value (line with the open circles) are shown.

**Figure 3.**
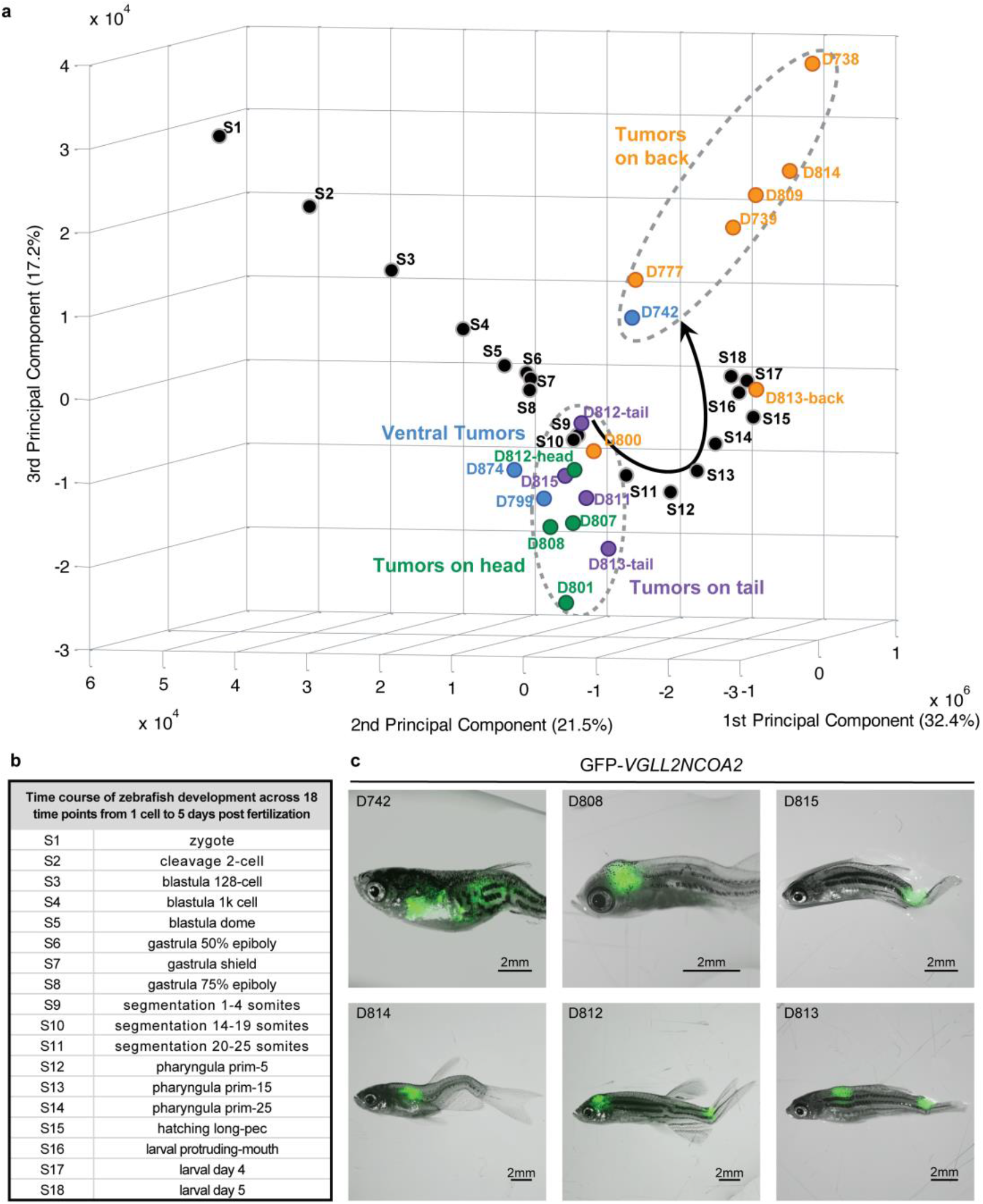
Gene expression profiles of *VGLL2-NCOA2* zebrafish tumors cluster with developmental timepoints. A) Principal component analysis of RNA sequencing data of zebrafish development time points from White et al 2017 ^27^ and *VGLL2-NCOA2* zebrafish tumors from this study. All FASTQ files were processed by the same computational pipeline to minimize computational batch effects. Colors indicate location of the tumor on the fish: back tumors in orange, ventral tumors in blue, head tumors in green and tail tumors in purple. In the PCA, PC1 describes the most variance, but is presented to appreciate differences in PC2 which best discriminates tumor cohorts. B) Embryonic stages labeled as per the Zebrafish Information Network (ZFIN). C) Brightfield and GFP overlaid images of a subset of the *VGLL2-NCOA2* tumors used in A.

We then performed RNA-sequencing (RNA-seq) on a cohort of zebrafish VGLL2-NCOA2 sarcomas to determine how well they transcriptionally recapitulate the human disease. To do this, we used a comparison of *VGLL2-NCOA2* tumors to another sarcoma model, *CIC-DUX4* Ewing like sarcoma. Both *VGLL2-NCOA2* and *CIC-DUX4* driven zebrafish tumors were compared to their human counterpart in an Agreement of Differential Expression analysis with a minimum of 5000 permutations (Figure 2C) ^4,24,26^. There were 2918 genes that were shared and differentially expressed between the zebrafish and human *VGLL2-NCOA2* tumor types as compared to *CIC-DUX4*. In Figure 2C, the cosine of the angle (cos) and difference of proportions (dop) statistics are both positive and significant, suggesting the zebrafish model recapitulates the human disease. This is supported by a Fisher’s exact test indicating a significant enrichment in shared differentially expressed genes between human and zebrafish *VGLL2-NCOA2* tumors as compared to *CIC-DUX4* tumors. Gene ontology analysis of this list of 2918 overlapping genes revealed multiple GO and KEGG terms related to skeletal muscle, differentiation, cellular proliferation, actin remodeling and developmental processes (Figure 2D). Overall, these data support implementing zebrafish tumor models to understand human *VGLL2-NCOA2* disease biology.

### VGLL2-NCOA2 leverages developmental genes and programs during tumor formation

Given the early onset of the tumors in the human and zebrafish cohort, and the possible link to development observed in the RNA-seq data, we hypothesized that the fusion oncogene was co-opting developmental genes to promote tumorigenesis. RNA-sequencing data from 18 zebrafish developmental timepoints from one-cell to five days post fertilization were analyzed alongside *VGLL2-NCOA2* zebrafish tumors, and were plotted using the first three principal components ^27^. This analysis revealed a large cohort of tumors clustered with the developmental timepoints of segmentation (Figure 3A; Stage S9-S11). This specific developmental timepoint is striking as it corresponds to somitic muscle development in the zebrafish (Figure 3B). The tumors that clustered with segmentation include the tumors on the head (green, Figure 3A), tail (purple, Figure 3A) and several of the ventral tumors (blue, Figure 3A). However, the tumors on the back (orange, Figure 3A) did not cluster with this group, instead forming a second cluster distant from the developmental trajectory.

To identify genes in *VGLL2-NCOA2* tumors that account for this clustering with segmentation, we performed a differential gene expression analysis from the RNA-seq data. First, the cohort of n=18 CMV-*VGLL2NCOA2* driven zebrafish tumors was compared to n=7 mature zebrafish skeletal muscle samples, identifying 385 differentially regulated genes. Next, n=3 samples of embryonic zebrafish during somitogenesis (1-4 somites, 14-19 somites, and 20-25 somites) were compared to mature zebrafish skeletal muscle; and 303 differentially regulated genes were found. Then, these 385 and 303 gene sets were intersected to identify 27 genes shared in both development and tumor datasets as differentially expressed when compared to mature skeletal muscle (Figure 4A). These 27 genes represent potential targets that are reactivated, inappropriately persist, or are never turned on in tumors. To understand the magnitude of this expression, the FPKM of these 27 genes was plotted, including genes downregulated in the tumors and developing muscle as compared to adult skeletal muscle (Figure 4B), and genes that were conversely upregulated in tumors and development, as compared to adult skeletal muscle (Figure 4C). Hierarchical clustering using Spearman Rank analysis on this 27 gene set revealed two main clusters that separated the adult muscle from the tumors and developmental samples (Figure 4D, Supplemental Figure 4A-B). In a further sub-grouping, the tumors along the back tend to cluster distinctly from the developmental samples and head and tail tumors, which is consistent with the sub-grouping observed in the PCA analysis (Figure 3A). Interestingly, although muscle regulatory factors were expressed in *VGLL2-NCOA2* tumors, there was not a significant difference in the expression level as compared to mature skeletal muscle or developmental samples (Supplemental Figure 3).

**Figure 4.**
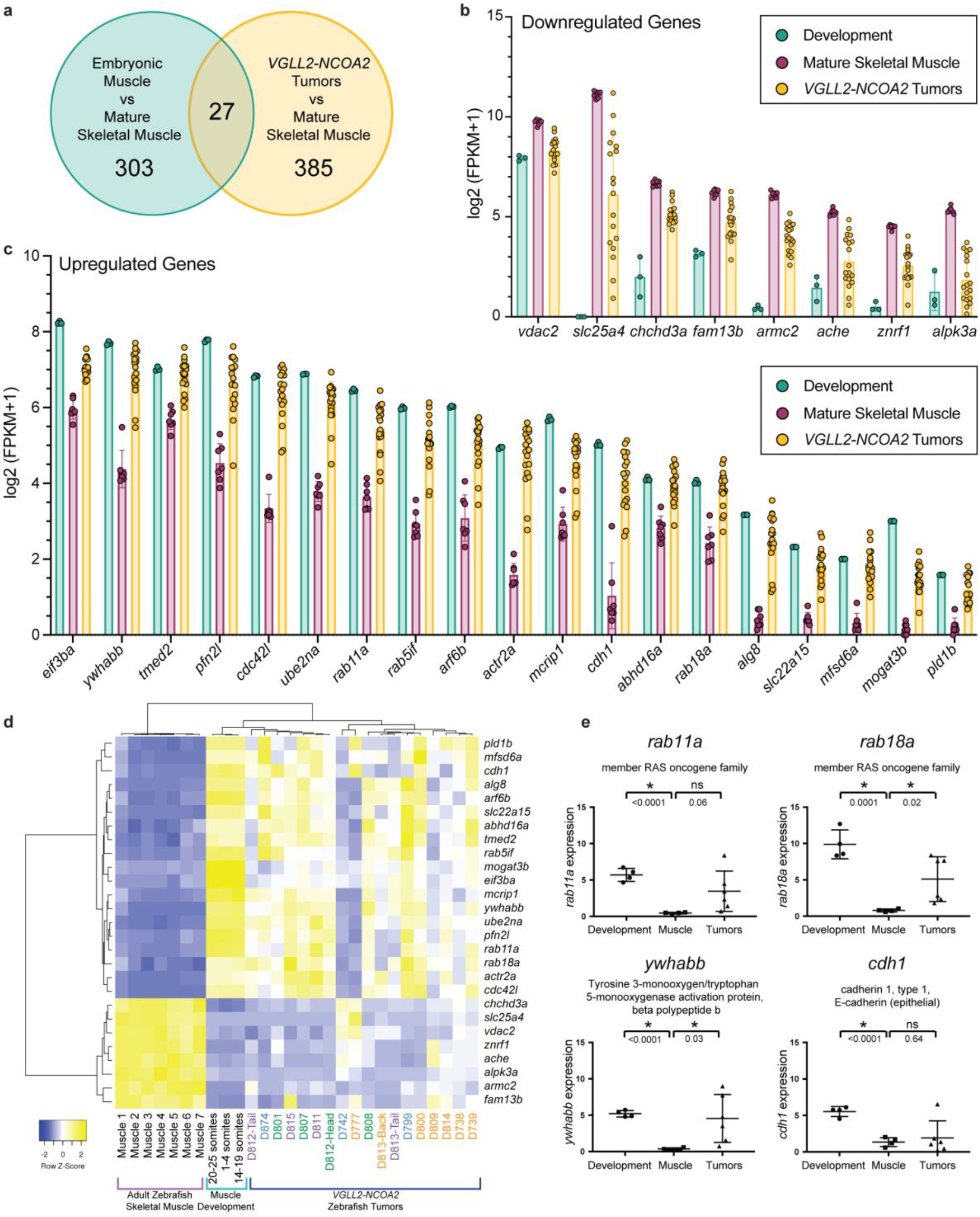
*VGLL2-NCOA2* reactivates developmental gene targets in tumors. RNA sequencing was performed on n=18 zebrafish *VGLL2-NCOA2* tumors, n=7 adult zebrafish back skeletal muscle samples and n=3 pooled samples of larval zebrafish from segmentation timepoints at 1-4 somites, 14-19 somites, and 20-25 somites (10.33, 16, and 19 hours post fertilization at 28°C, respectively). A) Venn diagram depicting the genes differentially regulated in the developmental timepoints and tumors as compared to mature skeletal muscle, with n=27 genes differentially regulated in both tumors and development. A FDR-adjusted *P* value cutoff of 0.01 was used for genes to be included in this analysis. B) Plot of FPKM values from n=8 genes downregulated in developmental timepoints and tumors as compared to mature skeletal muscle. C) Plot of FPKM values from n=19 genes upregulated in developmental timepoints and tumors as compared to mature skeletal muscle. D) Heat map and clustering analysis of the FPKM of the n=27 differentially expressed genes (from 4A-C) in *VGLL2-NCOA2* tumors, mature zebrafish skeletal muscle, and developmental timepoints. The samples are clustered by row and column using the average linkage clustering method and Spearman rank correlation for distance measurement method. Tumors are colored based on location on the fish: back tumors in orange, ventral tumors in blue, head tumors in green and tail tumors in purple. E) qRT-PCR of selected genes from A to evaluate using an independent cohort of n=6 zebrafish tumors, n=4 zebrafish mature skeletal muscle samples and n=4 pools of zebrafish embryos at a developmental timepoint of 20 hours post fertilization. Each datapoint is an individual tumor or normal tissue sample. The error bars represent the mean ± SD. *P* values were calculated using a two-tailed Student’s t-test.

We confirmed several of the developmental genes in this 27 gene set by qRT-PCR using independent cohorts of zebrafish tumors, development samples from 20 hours post fertilization and adult zebrafish skeletal muscle. This subset of tested genes included cadherin 1, ywhabb, and several members of the RAS superfamily (Figure 4E). Overall, this analysis identified genes whose expression in tumors more closely matched that in embryos rather than mature skeletal muscle, suggesting that *VGLL2-NCOA2* co-opts embryonic gene expression programs to promote tumor growth.

### Cross-species comparative oncology analysis identifies ARF6 as upregulated in zebrafish, mouse allograft, and human VGLL2-NCOA2 tumors

Gene ontology analysis of genes differentially expressed between *VGLL2-NCOA2* zebrafish tumors and mature skeletal muscle highlighted several GTPases and their functions (Figure 5A). One of the genes upregulated in both zebrafish development and *VGLL2-NCOA2* tumors was *arf6b*, a small GTPase involved in endocytosis and membrane receptor recycling and in actin reorganization (Figure 4D) ^28^. In zebrafish, the *arf6* gene is duplicated and is referred to as *arf6a* and *arf6b*. The zebrafish arf6a and arf6b protein sequences are identical and share 99% identity with the ARF6 human ortholog (Supplemental Figure 5). We performed qRT-PCR on both *arf6a* and *arf6b* to determine mRNA expression levels in an independent zebrafish tumor cohort, and found that both *arf6a/b* were elevated in development and tumor samples as compared to adult skeletal muscle (Figure 5B). We performed IHC for arf6 on serial sections of zebrafish *VGLL2-NCOA2* tumors and found that arf6a/b protein expression overlaid with tumor cells and human anti-NCOA2 positive cells (Figure 5C). These data suggest that arf6 is highly expressed in *VGLL2-NCOA2* established tumors and could be contributing to the disease process.

**Figure 5.**
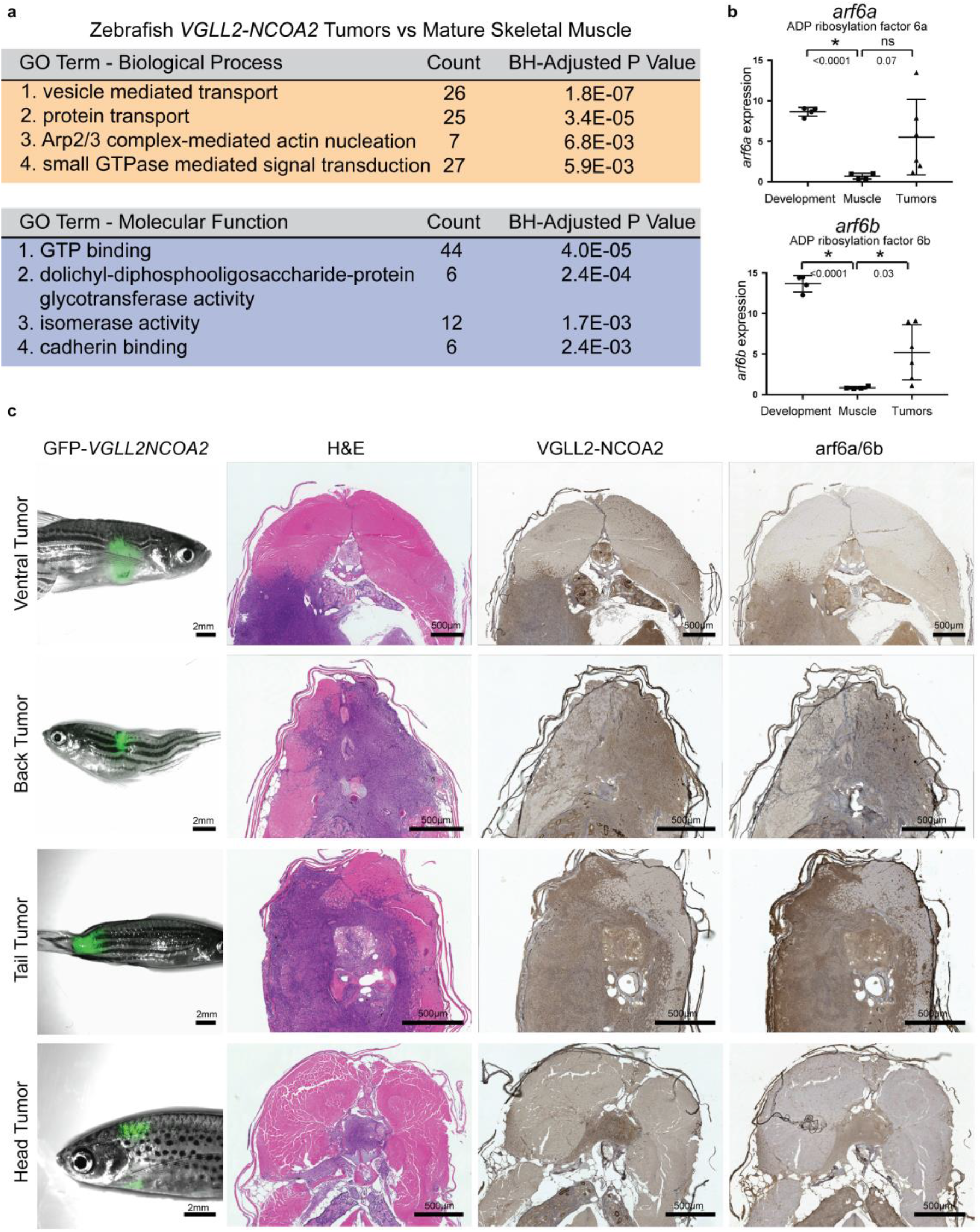
Zebrafish *VGLL2-NCOA2* tumors express arf6 protein. A) Gene ontology terms associated with differentially expressed genes in *VGLL2-NCOA2* zebrafish tumors versus mature zebrafish skeletal muscle. B) qRT-PCR validation of *arf6a* and *arf6b* levels using the samples from Figure 4E. C) Representative images of *VGLL2-NCOA2*zebrafish tumors shown as brightfield overlaid with GFP fluorescence, with serial transverse sections stained with hematoxylin and eosin, an anti-NCOA2 antibody and an anti-ARF6 antibody. All n=8 zebrafish tumors stained for ARF6 were positive.

To implement a cross species comparative oncology approach, we then generated a C2C12 mouse myoblast cell line that constitutively and stably overexpressed the human form of the *VGLL2-NCOA2* fusion that is used in our zebrafish models. We found that the fusion was highly expressed at the protein level even after passage and selection in culture (Figure 6A). Further, when either C2C12-pCDNA3.1 control or C2C12-*VGLL2NCOA2* cells were allografted into SCID or NUDE mice, cells overexpressing the *VGLL2-NCOA2* fusion generated aggressive tumors that rapidly grew with initial detection at eleven days post injection and termination of the experiment at 26-28 days post injection (Figure 6B-C). This indicates that the human form of the fusion-oncogene is transforming in multiple systems. We performed RNAseq on a sub-set of these generated C2C12-pCNDA3.1 control and C2C12-*VGLL2NCOA2* allografts, and found that *Arf6* was significantly overexpressed in the context of the *VGLL2-NCOA2* fusion-oncogene (Figure 6D). Next, we analyzed RNA-seq data from patient tumors, and performed differential expression analysis between human *VGLL2-NCOA2* tumors and human mature skeletal muscle and found that *ARF6* mRNA was modestly overexpressed in the human disease (Figure 7A). This observation may be translatable to many other sarcomas, with similar overexpression of *ARF6* as compared to mature skeletal muscle being observed in adult sarcomas in general, and in pediatric sarcomas, including Ewing sarcoma, osteosarcoma, and rhabdomyosarcoma. Notably, this did not hold true for clear cell sarcoma of kidney (Figure 7B).

**Figure 6.**
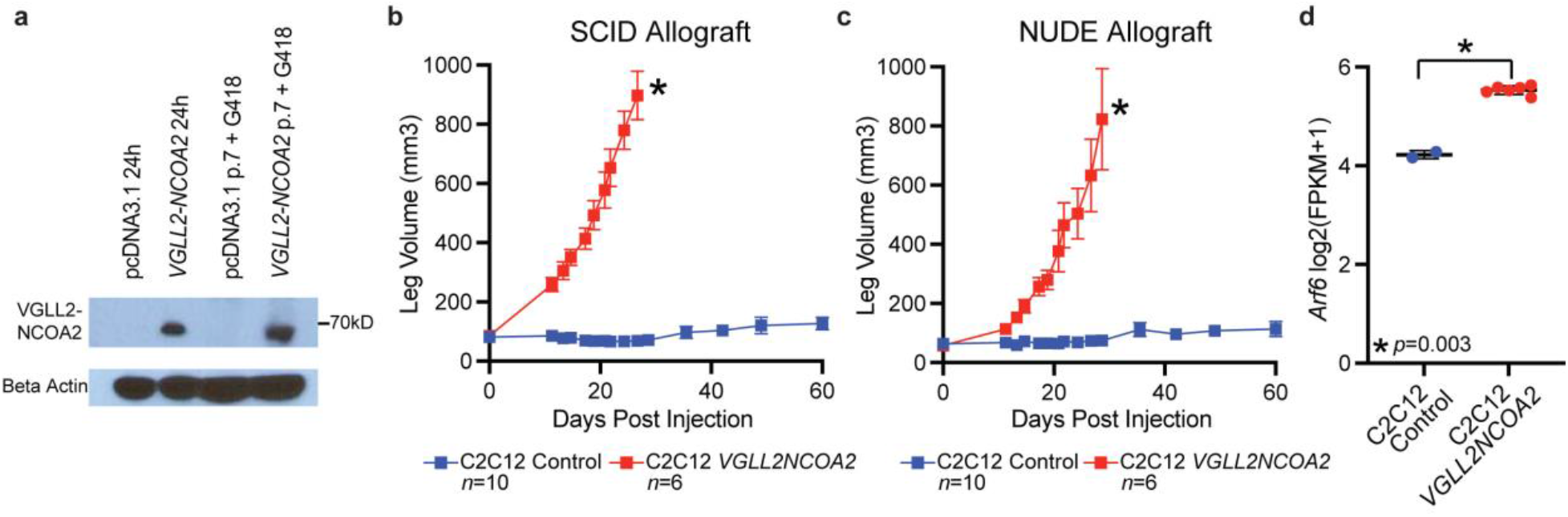
*VGLL2-NCOA2* is transforming in mouse myoblast allograft models. A) A C2C12 mouse myoblast human *VGLL2-NCOA2* transfected cell line expresses the VGLL2-NCOA2 fusion protein 24 hours post transfection and stably after passaging and selection in growth media with G418 as compared to a C2C12-pcDNA3.1 empty control. B) Allograft tumor volume of C2C12-pcDNA3.1 control and *C2C12-VGLL2NCOA2* after intramuscular injection into the leg of SCID mice. C2C12-pcDNA3.1 control was injected in n=10 allografts and C2C12-*VGLL2NCOA2* was injected in n=6 allografts. The error bars represent the mean tumor volume ±SEM. Error bars are not shown if it is within the boundaries of the symbol. Tumor volume at timepoints were compared using a Mann-Whitney U test corrected for multiple comparisons using the Benjamini, Krieger and Yekutieli method. * indicates *P* <0.0005. Every timepoint after zero is statistically significant. C) Allograft tumor volume of C2C12-pcDNA3.1 control and C2C12-*VGLL2NCOA2* after intramuscular injection into the leg of Swiss Nude mice. C2C12- pcDNA3.1 was injected in n=10 allografts and C2C12-VGLL2NCOA2 was injected in n=6 allografts. The error bars represent the mean tumor volume ± SEM. Error bars are not shown if it is within the boundaries of the symbol. Tumor volume at timepoints were compared using a Mann-Whitney U test corrected for multiple comparisons using the Benjamini, Krieger and Yekutieli method. * indicates *P*<0.0005. Every timepoint after zero is statistically significant. D) *Arf6* mRNA levels from RNAseq of allografts from Swiss Nude mice, including two C2C12- pcDNA3.1 controls and six C2C12-VGLL2NCOA2 tumors. *Arf6* mRNA expression was compared using a Welch’s t-test.

**Figure 7.**
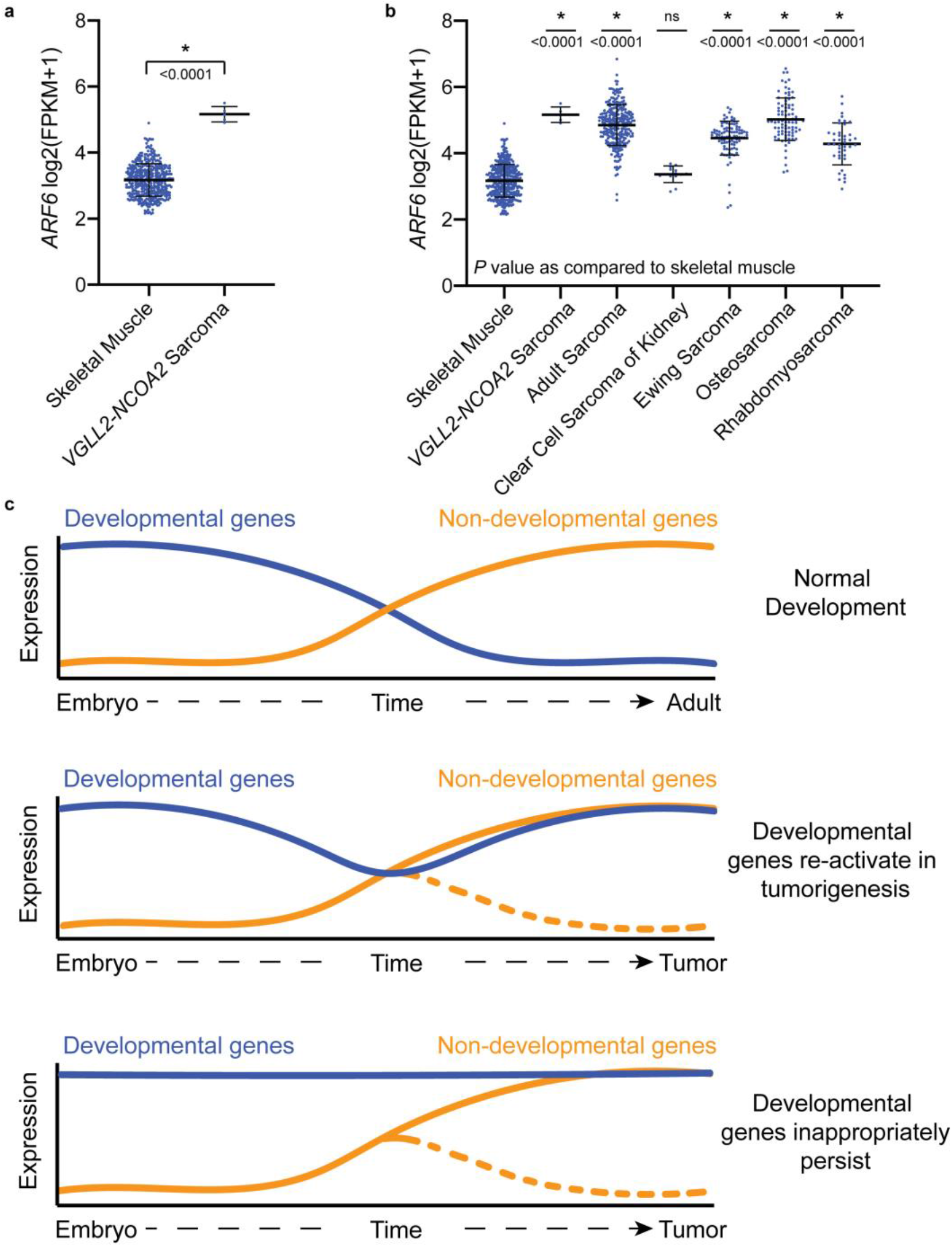
*ARF6* expression in patient sarcomas and a model for *VGLL2-NCOA2* tumorigenesis. A) *ARF6* expression determined by RNA-seq in skeletal muscle (n=396) and *VGLL2-NCOA2* tumors from Watson et al 2018 (n=5; one is a primary and recurrent from the same patient) ^4^. Each datapoint is an individual tumor or normal tissue sample. The error bars represent the mean ± SD. *P* values were calculated using the Mann-Whitney U Test. B) *ARF6* expression from (A) and compared to additional sarcoma tumor samples; including, adult sarcoma from TCGA (n=264), clear cell sarcoma of kidney (n=13), Ewing sarcoma (n=95), osteosarcoma (n=87), and rhabdomyosarcoma (n=42). Each datapoint is an individual tumor or normal tissue sample. The error bars represent the mean ± SD. P values were calculated using the Mann- Whitney U Test and are a comparison of each tumor cohort to skeletal muscle. C) Models of *VGLL2-NCOA2* tumorigenesis. Development is a tightly regulated process where genes typically have discrete developmental roles and are rarely repurposed for use in adult or mature tissues. Non-developmental genes are expressed to maintain homeostasis. A potential mechanism for *VGLL2-NCOA2* oncogenic activity is to leverage the unique properties of developmental genes for rapid growth and inhibition of differentiation, and to either re-activate them, or allow them to inappropriately persist, facilitating the tumorigenic process.

Overall, development is a tightly controlled and orchestrated process involving appropriate gene activation and inactivation to dictate migration, cell division, and structure formation. Our data suggest that *VGLL2-NCOA2* is an oncogene which potentially leverages developmental programs in order to promote sarcomagenesis. There are two distinct but not mutually exclusive possibilities for the mechanism: one in which during maturation developmental genes continuously decrease in expression, and are then are re-activated either at the onset or during tumorigenesis. Alternatively, developmental genes may never decrease in expression and instead inappropriately persist throughout the course of tumorigenesis (Figure 7C). Further studies will be required to delineate the mechanisms by which *VGLL2-NCOA2* mediates these changes in gene expression, and their role in tumorigenesis.

## Discussion

Each sarcoma gene fusion produces a slightly different disease that warrants the development of a genetic animal model system to understand the biology and to identify tractable therapeutic targets. This is difficult to accomplish. Often, the sarcoma cell of origin is unknown making screening putative cellular origins in mouse models costly and burdensome. Zebrafish complement mouse models as a timely approach to make significant contributions to understanding rare genetic diseases and pediatric cancer. Zebrafish capture a relevant developmental context and have conserved genetics and molecular signaling pathways ^29–31^. By applying these models, one can complement clinical sequencing efforts and define the biology of newly discovered fusion-oncogenes to determine the most significant features of the disease. Previously, we described a genetic approach focused on *PAX3-FOXO1* rhabdomyosarcoma and *CIC-DUX4* sarcoma transgenic zebrafish modeling ^23,24^. We have now broadened our approach to incorporate newly discovered rhabdomyosarcoma fusion-oncogenes. This modular platform can be implemented for any rhabdomyosarcoma fusion identified from clinical sequencing efforts that has yet to be studied in a laboratory setting.

Here, we applied this strategy to demonstrate that the new putative cancer gene fusion, *VGLL2-NCOA2*, is an oncogene and generates sarcoma in genetic zebrafish and mouse allograft systems. When the human form of the fusion is integrated into the zebrafish genome, zebrafish develop tumors that are histologically and molecularly consistent with the human disease. Introduction of the VGLL2-NCOA2 fusion into C2C12 mouse myoblasts and allografts of these models result in aggressive tumors by three weeks post injection. These data indicate the fusion alone is sufficient for transformation and does not require cooperating events. This highlights the capacity of the zebrafish system to faithfully capture distinct sarcoma subtypes based on the driving oncogene. For example, zebrafish *VGLL2-NCOA2* tumors present at a much earlier age, have distinct histology and unique gene expression profiles as compared to our *PAX3-FOXO1* zebrafish tumor model ^23^. This is consistent with the human disease, in which patients with *VGLL2-NCOA2* tumors present between 0-1 year with distinct histology from *PAX3-FOXO1* driven rhabdomyosarcoma where they typically present during adolescence. Our data underscores the importance of creating and studying animal models of rare genetic diseases to determine biological differences and similarities in pediatric sarcomas.

In infantile rhabdomyosarcoma there are other fusion partners for *VGLL2*, including *CITED2* ^4,8,18,20^. However, transcriptional clustering analysis indicates *VGLL2*-fused tumors have shared signatures, suggesting that the most important disease driver is *VGLL2*, a transcriptional co-activator, and its regulatory elements ^4,20^. Insight into normal VGLL2 expression and its targets may help understand its activity in the fusion. *VGLL2* is highly expressed during mouse embryonic development in the somitic myotome and pharyngeal pouch, but in adult tissues its expression is restricted to skeletal muscle ^32^. In human development, *VGLL2* is solely expressed in fetal skeletal muscle, and is absent from cardiac, liver, thymus, or brain ^32^. This specific expression and interaction partners could explain the immature muscle features of the disease. A *Vgll2* knock-out mouse has helped elucidate its role in skeletal muscle development, and *Vgll2* knock-out mice exhibit defects in skeletal muscle fiber type composition and exercise exhaustion ^33,34^. This model has confirmed direct interaction with Vgll2 and TEAD1/4 by co-immunoprecipitation in embryonic skeletal muscle ^33^. Intriguingly, *TEAD1-NCOA2* fusions are another defining genetic event in congenital sarcoma with a similar clinical presentation ^18,19^. Perhaps, VGLL2 is responsible for regulating targets through TEAD1/4 proteins for precise developmental control, and this could be co-opted or deregulated by the *VGLL2-NCOA2* fusion.

Transcriptional analysis of *VGLL2-NCOA2* zebrafish tumors identified a subset that clustered with somitogenesis in the developing fish, distinct from zebrafish mature skeletal muscle. These results suggest that tumor programs either reactivate developmental genes or that developmental genes inappropriately persist. Our differential gene expression analysis highlighted small GTPases. Specifically, of these GTPases, arf6 had increased protein expression in zebrafish tumor cells and was completely absent in adult skeletal muscle. The sufficiency of the *VGLL2-NCOA2* fusion and overexpression of *Arf6* in the contest of the fusion was confirmed in C2C12 mouse allograft models. In the human disease, *ARF6* mRNA is overexpressed in *VGLL2-NCOA2* patient tumors as compared to mature skeletal muscle. More broadly, in a panel of pediatric and adult sarcomas, *ARF6* is also over-expressed as compared to mature skeletal muscle suggesting that ARF6 could be a therapeutic target in many sarcoma contexts. Perhaps this has been overlooked because *ARF6* mRNA levels do not always correlate with ARF6 protein expression (Figure 5B-C) ^35^, and ARF6 is not a defining feature of any sarcoma subtype but represents a shared feature of the disease.

ARF6 is a highly conserved mediator of developmental processes, myogenesis, and carcinogenesis, all of which could contribute to its function in *VGLL2-NCOA2* tumors. Studies in sea urchins suggest that *Arf6* knock-down impairs gastrulation and alters directed migration of primary mesenchyme cells ^36^. This is complemented from data in *Arf6* null mice, which are embryonic lethal indicating a critical developmental role ^37^. ARF6 is also a core member of the complex required for myoblast fusion and muscle development ^38^. Impaired muscle differentiation is a hallmark of rhabdomyosarcoma, and these processes are aberrant in *VGLL2-NCOA2* tumors, which express the early markers of myogenesis but appear histologically undifferentiated. ARF6 activation or overexpression has a role in other cancer contexts and promotes proliferation, invasion, and metastasis or predicts poor prognosis in breast cancer, melanoma, lung cancer, and other solid tumors ^35,39–43^. A remaining question is the contribution of ARF6 to tumor initiation versus maintenance (or both), which is not clear, and will be a focus of future lines of research. ARF6 has also been proposed as a therapeutic opportunity ^44^. In uveal melanoma, an ARF6 inhibitor, NAV-2729 inhibited tumor growth in xenograft models of the disease ^40^. Although ARF6 is broadly expressed, the tolerability of NAV-2729 in these studies indicates that there may be a therapeutic window. Whether this is true in *VGLL2-NCOA2* driven rhabdomyosarcoma, or sarcoma in general, remains to be seen.

Overall, our cross species comparative oncology approach developed the first animal and allograft model of *VGLL2-NCOA2* fusion-driven sarcoma and have identified a potentially druggable target, ARF6. Our findings also highlight how studying rare disease can illuminate molecular mechanisms or systems that are more broadly applicable. We have now presented a roadmap for building new fusion-driven animal models to define biology, identify therapeutic opportunities, and intersect data to determine conserved sarcoma disease drivers.

## Methods and Materials

### Zebrafish husbandry

*Danio rerio* were maintained in an Aquaneering aquatics facility according to industry standards. Vertebrate animal work is accredited by AALAC and overseen by the UT Southwestern Medical Center or Nationwide Children’s Hospital IACUC committee and Animal Resources Core. Wildtype lines used in this study were AB, WIK, TL and were obtained from the Zebrafish International Resource Center (ZIRC). AB/TL were bred in our facility by crossing AB and TL. The *p53* mutant line, *tp53^M214K^*, was a kind gift from Tom Look ^45^.

### Plasmids and cloning

*VGLL2-NCOA2* coding sequence was cloned out of a patient tumor. In the fusion used to develop the zebrafish model, exons 1 and 2 of *VGLL2* is fused to exon 14 through 23 of *NCOA2*. Sanger sequencing revealed that serine at position 544 of the resulting fusion protein had the nucleotide sequence TCT, a silent mutation compared to the human reference sequence of TCC. The fusion was cloned into the Gateway expression system by adding 5’ and 3’ ATT sites (attb2r/attb3) using a high-fidelity polymerase. Purified PCR product was then cloned into a 3’ entry clone as described in ^46^. The Tol2 kit components: destination vector pDestTol2pA2, 3’ SV40 late polyA signal construct, beta actin promoter, cmv promoter, and multiple cloning site were used to generate constructs for expression in zebrafish ^47^. The ubi promoter was a kind gift from Len Zon (Addgene #27320) ^48^, and the unc503 promoter from Peter Currie (Addgene #27320) ^49^. Middle entry beta globin intron and splice acceptor was from Koichi Kawakami ^50^. The GFP-viral2A sequence were a gift from Steven Leach ^51^ and were sub-cloned into a middle entry Gateway expression system ^23^. Tol2 mRNA was synthesized in vitro from pCS2FA-transposase which is from Koichi Kawakami ^25^. The CMV-GFP2A-pA construct was generated as described in Kendall et al., 2018 ^23^. The zebrafish expression constructs generated for this study include: CMV-GFP2A-*VGLL2NCOA2*, BetaActin-GFP2A-*VGLL2NCOA2*, ubi-GFP2A-*VGLL2NCOA2*, unc503-GFP2A-*VGLL2NCOA2*, MCS-beta-globin-SpliceAcceptor-GFP2A-*VGLL2NCOA2*. For expression in C2C12 cells, the VGLL2-NCOA2 fusion was sub-cloned into pcDNA3.1 using primers that added BamH1 and Xho1 digest sites to the 5’ and 3’ ends of the transcript, respectively (Supplemental Table 3).

### Zebrafish embryo injections

Zebrafish embryos were injected at the single-cell stage with an injection mix containing 50 ng/μL of Tol2 transposon mRNA, 50 ng/μL of the *VGLL2-NCOA2* DNA construct and equimolar amounts of control constructs, 0.1% phenol red and 3x Danieau’s buffer.

### Zebrafish tumor collection and processing

Zebrafish with tumors were euthanized in a 0.1% Tricaine-S solution and screened using the Nikon SMZ25 fluorescent stereoscope to detect GFP expression. Tumors were resected and flash frozen in liquid nitrogen. Total RNA isolation was performed on frozen tissue using the RNeasy Micro Kit (Qiagen #74004). The remaining zebrafish was fixed in 4% paraformaldehyde in PBS for 48 hours shaking at 4°C, subsequently decalcified in 0.5M EDTA for 5 days at room temperature and mounted in paraffin blocks for microtome sectioning. Sections were taken at 5-10 micron intervals. De-paraffinized slides were stained with Hematoxylin and eosin as described in Kendall and Amatruda, 2016 ^46^.

### Zebrafish tumor incidence

All zebrafish that survived past thirty days were counted for tumor incidence curves. Zebrafish with no GFP fluorescence were counted as negative for transgene-dependent tumor formation. Zebrafish with GFP fluorescence were processed as described earlier and tumor formation was confirmed by visual review by a pathologist of hematoxylin and eosin stained slides. Fish tumors were classified by location (Figure 1C): tumors anterior to the gill were considered head tumors, and tumors posterior to the largest part of the ventral fin were considered tail tumors. Tumors between these two boundaries were divided into back tumors and ventral tumors at the midcoronal plane. Tumors spanning more than one area were classified by the area in which the tumor predominately presented.

### Generation of C2C12-VGLL2NCOA2 cell line

C2C12 cells were obtained from American Type Culture Collection (#CRL-1722) and were cultured in growth media consisting of 20% fetal bovine serum in DMEM (GE healthcare # SH30022.01). Cells were transfected with pcDNA3.1 empty vector or pcDNA3.1 containing the human coding sequence of the VGLL2-NCOA2 fusion using the AMAXA cell line Nucleofection kit V (Lonza #VCA-1003;) and were selected in growth media containing 1μg/ml concentration of G418 (ThermoFisher #10131035). Cells expressed the VGLL2-NCOA2 fusion at 24 hours post transfection and stably after passage in culture under G148 selection.

### Western blot

C2C12 cell lysate was generated using 5 million cells in RIPA lysis buffer with protease/phosphatase inhibitors (Roche #11836145001 and #04906837001), and 20μg of protein was loaded on a denaturing SDS page gel for detection of the VGLL2-NCOA2 fusion. Western blot was performed using primary antibodies against human NCOA2 at 1:5000 dilution (Abcam #Ab10508,) and anti beta actin at 1:10000 dilution (Sigma #A5316), and were visualized using monoclonal anti-Rabbit (1:3000) secondary, and chemiluminescence on film.

### Allograft mouse experiments

Animal care and use for this study were performed in accordance with the recommendations of the European Community (2010/63/UE) for the care and use of laboratory animals.

Experimental procedures were specifically approved by the ethics committee of the Institut Curie CEEA-IC #118 (Authorization APAFIS#11206-2017090816044613-v2 given by National Authority) in compliance with the international guidelines. C2C12-pcDNA3.1 control and C2C12-pcDNA3.1-VGLL2NCOA2 cell lines were utilized in xenograft experiments. Two million cells were resuspended in 50μL of sterile 1X PBS and injected intramuscularly into the leg of SCID (Charles River Laboratories CB17/Icr-*Prkdc^scid^*/IcrIcoCrl) or Swiss Nude (Charles River Laboratories Crl:NU(Ico)-*Foxn1^nu^*) mice under anesthesia. All injected mice were female and, except for one Nude mouse at 136 days of age, were 46 days old at the time of injection. Tumors were detected eleven days after injection, after which the height and width of the injected leg were measured using calipers to calculate the volume until 60 days post injection for C2C12 controls, and until 26 or 28 days post injection for C2C12-VGLL2NCOA2 injected mice (initial leg volume was 50-100mm^3^). At these later timepoints mice were euthanized due to tumor burden (C2C12-VGLL2NCOA2) or due to the end of the experiment (C2C12-pcDNA3.1 control).

### Immunohistochemistry staining

Slides were baked in a 60°C oven for 1 hour, deparaffinized with xylene/Histo-Clear for two incubations of 10 min, rehydrated in two 3 min washes in 100% Ethanol, 95% Ethanol and then diH_2_O. Antigen retrieval was performed in Trilogy reagent for 15 min in a pressure cooker. The slides were incubated with 3% peroxidase H_2_O_2_ for 30 min, washed with water for 1 min, blocked with 1% BSA/1x PBST for 1 hour. Primary antibody incubation was done at 1:100 dilution at 4°C overnight. Primary antibodies used were anti-NCOA2 (Novus BIO NB100-1756) and anti-ARF6 (Abcam ab77581). Slides were washed with PBST 4 times for 5 min each, and HRP secondary conjugate (BioRad 172-1019) was applied at 1:300 dilution for 1 hour at room temperature. Washes were performed as above, and DAB solution (Sigma #D5905) with 0.03% H_2_O_2_ applied for 5 min. Slides were counterstained with hematoxylin for 5 min and dehydrated in 95% Ethanol, 100% Ethanol, and Xylene/Histo-Clear each for two incubations of 1 min each. Slides were imaged on a Keyence BZ-X710 microscope.

### RNA isolation, RT-PCR and qRT-PCR

RNA was isolated from zebrafish tumors, normal tissue, or embryos using the QIAGEN RNeasy micro or mini-kit depending on the tumor size. For skeletal muscle, an additional proteinase K digestion step was included. 500ng of RNA isolated from zebrafish tumors, normal tissue or n=30 embryos at 20 hours post fertilization was used as input to the RT^2^ HT First Strand Kit (Qiagen #330411) to reverse transcribe cDNA. Standard PCRs were run using Taq Polymerase (NEB) using primers provided in Supplemental Table 3. qRT-PCR was performed on the BioRad CFX384 using standard cycling conditions and the 2X BioRad Master Mix. Data were analyzed in CFX Maestro Software and are plotted as the values relative to zero as normalized to two input controls, *rplp13a* and *gapdh*.

### RNA sequencing and analysis

Approximately 2μg of RNA was used for each zebrafish sample run. RNA-seq for D738, D739, D742, D777, D799, D800, D801, D807, D808, D813, Muscle 2 and Muscle 3 was done at the Institut Curie. Total RNA was isolated from Nude C2C12-pcDNA3.1 control or *C2C12-VGLL2NCOA2* allografts using Trizol, and approximately 1 ug was used for RNA-seq at the Institut Curie. D809, D811, D812, D814, D815, D874, and additional five mature skeletal muscle samples (D948, D949, D950, D951, D952) were sent to DNALinks for mRNA sequencing with poly-A RNA enrichment (2×75bp run, 70M reads of data generation). All tumors were generated in an AB/TL wildtype background, except for D738 and D742 which are in the *tp53^M214K^* homozygous mutant background ^45^. Additional sample details are in Supplemental Table 4. Trim Galore (https://www.bioinformatics.babraham.ac.uk/projects/trim_galore/) was used for quality and adapter trimming. The qualities of RNA-sequencing libraries were estimated by mapping the reads onto zebrafish reference dataset (Genome Reference Consortium Zebrafish Build 11, GRCz11) using Bowtie (v2.3.4.3) ^52^. STAR (v2.7.2b) ^53^ was employed to align the reads onto the zebrafish genome GRCz11, SAMtools (v1.9) ^54^ was employed to sort the alignments, and HTSeq Python package ^55^ was employed to count reads per gene. edgeR R Bioconductor package ^56–58^ was used to normalize read counts and identify differentially expressed (DE) genes. The enrichment of DE genes to pathways and GOs were calculated by Fisher’s exact test in R statistical package ^59^. Differentially expressed genes were determined using cut-offs of fold changes > 1.5 and an FDR-adjusted p-value of < 0.05.

### RNA-seq comparison of zebrafish and human *VGLL2-NCOA2* tumors

Agreement of Differential Expression –AGDEX- ^26^ analysis was performed using RNA-seq data from our *CIC-DUX4* zebrafish model as compared to human *CIC-DUX4* sarcomas, and from *VGLL2-NCOA2* zebrafish model as compared to human *VGLL2-NCOA2* sarcomas ^4,24^. Gene expression values for both RNAseq series were assessed using Kallisto 0.46.1 ^60^ using indexes of the Homo sapiens GRCh38p4 genome or Danio rerio GRCz11 genome. Mapping of the genes between the two species was done using BioMart tool from the Ensembl website (https://www.ensembl.org/biomart/martview). AGDEX analysis was performed with a minimum number of permutations set to 5000.

### RNA-seq comparison of human *VGLL2-NCOA2* tumors, and other sarcomas, with mature skeletal muscle

RNA-seq data from *VGLL2-NCOA2* tumors is from Watson et al 2018 ^4^. RNA-seq data for n=264 adult sarcomas is from TCGA and represents dedifferentiated liposarcoma, leiomyosarcoma, undifferentiated pleomorphic sarcoma, myxofibrosarcoma, malignant peripheral nerve sheath tumor, and synovial sarcoma ^61^. RNAseq data from n=96 Ewing sarcoma tumors is from Crompton et al 2014 ^62^, RNAseq data from n=87 Osteosarcoma (phs000468) and n=13 Clear Cell Sarcoma of the Kidney (phs000466) are based upon data generated by the Therapeutically Applicable Research to Generate Effective Treatments (TARGET) (https://ocg.cancer.gov/programs/target) initiative, phs000468 and phs000466. The data used for this analysis are available at https://portal.gdc.cancer.gov/projects. RNA-seq data from n=42 fusion-positive and fusion-negative rhabdomyosarcoma is from Xu et al 2018 ^63^. RNA-seq data from n=396 adult skeletal muscle samples are from the Genotype-Tissue Expression Project (GTEx). The Genotype-Tissue Expression (GTEx) Project was supported by the Common Fund of the Office of the Director of the National Institutes of Health, and by NCI, NHGRI, NHLBI, NIDA, NIMH, and NINDS. The data used for the analyses described in this manuscript were obtained from the GTEx Portal on March 2018. The same computational analysis steps based on hg19 human reference genome data, as detailed in the RNA-seq and analysis section, were applied to process these tumor and muscle datasets side by side to minimize computational batch effects. In addition, we also applied ComBat to remove batch effects. HTSeq Python package ^55^ was employed to count reads per gene. edgeR R Bioconductor package ^56–58^ was used to normalize read counts and calculate FPKM values.

### PCA comparing zebrafish tumors to developmental stages

Zebrafish developmental RNAseq data were from White et al 20 1 7 ^27^. Zebrafish tumor and skeletal muscle sample RNAseq data were generated from this study. To minimize the computational batch effect, FASTQ files from White et al 2017 and this study were processed with the same pipeline, as detailed in the RNA sequencing and analysis section side by side to minimize computational batch effects from multiple data resources. We then performed the Principal Component Analysis based on princomp package in R.

### Heatmap Generation

The heatmap of FPKM values from RNA sequencing was generated using heatmapper.ca ^64^. The clustering method used was average linkage and Spearman rank correlation utilized for distance measurement method. Clustering was applied to rows and columns, and each row was scaled separately. Alternative distance measurement methods using Kendall’s Tau and Pearson were generated in the same manner.

### Gene Ontology Analysis

Genes with statistically significant expression changes were analyzed using DAVID (https://david.ncifcrf.gov/) ^65,66^ to identify enriched Gene Ontology (GO) terms (http://www.geneontology.org/). In Figure 5, the direct biological process (top) and molecular function (bottom) terms are listed for genes differentially regulated in zebrafish VGLL2-NCOA2 tumors versus skeletal muscle (p<0.01; FDR adjusted).

### Statistics

Analysis of survival curves, tumor incidence and onset curves, qRT-PCR analysis, and Fisher’s exact test was performed in Prism 8.1.1 (LaJolla, CA). Statistical tests in differences in tumor onset were performed using Log-Rank analysis. All other calculations were performed in Microsoft Excel Version 16.16.11. Sample sizes are provided in figures or figure legends.

## Acknowledgements

We thank the UT Southwestern Histopathology core for excellent services and for their expertise, and the Nationwide Children’s Hospital Animal Resources Core for their exceptional zebrafish husbandry. G.C.K is supported by an Alex’s Lemonade Stand Foundation “A” Award and a V Foundation for Cancer Research V Scholar Grant. J.F.A. was supported by grant RP120685-P3 from the Cancer Prevention and Research Institute of Texas (CPRIT) and by Curing Kids Cancer. L.X. is supported by Rally Foundation, Cancer Center Support Grant P30 CA142543 and RP180805 from CPRIT. F.T. is supported by the Institut National de la Recherche Médiacale (INSERM). S.W. was supported by a grant from the Fondation Nuovo-Soldati pour la recherche en cancérologie.

## Author Contributions

S.W, F.T., O.D., J.F.A and G.C.K conceived and supervised the study. F.T. provided computational analysis, insight, and expertise of human and zebrafish RNAseq data and their similarity in Figure 2. L.X. provided computational analysis, insight, and expertise into developmental studies and human and zebrafish RNAseq data in Figures 3 and 7. D.R provided pathological assessment. S.W., W.M., D.S., C.M.L, G.C.K performed experiments. C.M.L and G.C.K. analyzed experimental data. C.M.L, J.F.A., and G.C.K drafted the manuscript and figures. All authors reviewed and edited the final manuscript.

## References

1. Mitelman, F., B, J. & Mertens, F. Mitelman Database of Chromosome Aberrations and Gene Fusions in Cancer, <https://mitelmandatabase.isb-cgc.org> (2021).

2. Mitelman, F., Johansson, B. & Mertens, F. The impact of translocations and gene fusions on cancer causation. Nat Rev Cancer 7, 233–245, doi:10.1038/nrc2091 (2007).

3. Mertens, F., Antonescu, C. R. & Mitelman, F. Gene fusions in soft tissue tumors: Recurrent and overlapping pathogenetic themes. Genes Chromosomes Cancer 55, 291–310, doi:10.1002/gcc.22335 (2016).

4. Watson, S. et al. Transcriptomic definition of molecular subgroups of small round cell sarcomas. J Pathol 245, 29–40, doi:10.1002/path.5053 (2018).

5. Grobner, S. N. et al. The landscape of genomic alterations across childhood cancers. Nature 555, 321–327, doi:10.1038/nature25480 (2018).

6. Tirode, F. et al. Genomic landscape of Ewing sarcoma defines an aggressive subtype with co-association of STAG2 and TP53 mutations. Cancer Discov 4, 1342–1353, doi:10.1158/2159-8290.CD-14-0622 (2014).

7. Shern, J. F. et al. Comprehensive genomic analysis of rhabdomyosarcoma reveals a landscape of alterations affecting a common genetic axis in fusion-positive and fusion-negative tumors. Cancer Discov 4, 216–231, doi:10.1158/2159-8290.CD-13-0639 (2014).

8. Alaggio, R. et al. A Molecular Study of Pediatric Spindle and Sclerosing Rhabdomyosarcoma: Identification of Novel and Recurrent VGLL2-related Fusions in Infantile Cases. Am J Surg Pathol 40, 224–235, doi:10.1097/PAS.0000000000000538 (2016).

9. Gunther, S., Mielcarek, M., Kruger, M. & Braun, T. VITO-1 is an essential cofactor of TEF1-dependent muscle-specific gene regulation. Nucleic Acids Res 32, 791–802, doi:10.1093/nar/gkh248 (2004).

10. Maeda, T., Chapman, D. L. & Stewart, A. F. Mammalian vestigial-like 2, a cofactor of TEF-1 and MEF2 transcription factors that promotes skeletal muscle differentiation. J Biol Chem 277, 48889–48898, doi:10.1074/jbc.M206858200 (2002).

11. Pobbati, A. V. & Hong, W. Emerging roles of TEAD transcription factors and its coactivators in cancers. Cancer Biol Ther 14, 390–398, doi:10.4161/cbt.23788 (2013).

12. Johnson, C. W. et al. Vgll2a is required for neural crest cell survival during zebrafish craniofacial development. Dev Biol 357, 269–281, doi:10.1016/j.ydbio.2011.06.034 (2011).

13. Wang, L. et al. Identification of a novel, recurrent HEY1-NCOA2 fusion in mesenchymal chondrosarcoma based on a genome-wide screen of exon-level expression data. Genes Chromosomes Cancer 51, 127–139, doi:10.1002/gcc.20937 (2012).

14. Sumegi, J. et al. Recurrent t(2;2) and t(2;8) translocations in rhabdomyosarcoma without the canonical PAX-FOXO1 fuse PAX3 to members of the nuclear receptor transcriptional coactivator family. Genes Chromosomes Cancer 49, 224–236, doi:10.1002/gcc.20731 (2010).

15. Argani, P. et al. Novel MEIS1-NCOA2 Gene Fusions Define a Distinct Primitive Spindle Cell Sarcoma of the Kidney. Am J Surg Pathol 42, 1562–1570, doi:10.1097/PAS.0000000000001140 (2018).

16. Kao, Y. C. et al. Recurrent MEIS1-NCOA2/1 fusions in a subset of low-grade spindle cell sarcomas frequently involving the genitourinary and gynecologic tracts. Mod Pathol 34, 1203–1212, doi:10.1038/s41379-021-00744-7 (2021).

17. Agaram, N. P. et al. Expanding the Spectrum of Intraosseous Rhabdomyosarcoma: Correlation Between 2 Distinct Gene Fusions and Phenotype. Am J Surg Pathol 43, 695–702, doi:10.1097/PAS.0000000000001227 (2019).

18. Mosquera, J. M. et al. Recurrent NCOA2 gene rearrangements in congenital/infantile spindle cell rhabdomyosarcoma. Genes Chromosomes Cancer 52, 538–550, doi:10.1002/gcc.22050 (2013).

19. Whittle, S. B. et al. Congenital spindle cell rhabdomyosarcoma. Pediatr Blood Cancer 66, e27935, doi:10.1002/pbc.27935 (2019).

20. Butel, T. et al. Integrative clinical and biopathology analyses to understand the clinical heterogeneity of infantile rhabdomyosarcoma: A report from the French MMT committee. Cancer Med 9, 2698–2709, doi:10.1002/cam4.2713 (2020).

21. Anderson, W. J. & Doyle, L. A. Updates from the 2020 World Health Organization Classification of Soft Tissue and Bone Tumours. Histopathology 78, 644–657, doi:10.1111/his.14265 (2021).

22. Cyrta, J. et al. Infantile Rhabdomyosarcomas With VGLL2 Rearrangement Are Not Always an Indolent Disease: A Study of 4 Aggressive Cases With Clinical, Pathologic, Molecular, and Radiologic Findings. The American Journal of Surgical Pathology Publish Ahead of Print, doi:10.1097/pas.0000000000001702 (2021).

23. Kendall, G. C. et al. PAX3-FOXO1 transgenic zebrafish models identify HES3 as a mediator of rhabdomyosarcoma tumorigenesis. Elife 7, doi:10.7554/eLife.33800 (2018).

24. Watson, S. et al. CIC-DUX4 expression drives the development of small round cell sarcoma in transgenic zebrafish: a new model revealing a role for ETV4 in CIC-mediated sarcomagenesis. bioRxiv, 517722, doi:10.1101/517722 (2019).

25. Urasaki, A., Morvan, G. & Kawakami, K. Functional dissection of the Tol2 transposable element identified the minimal cis-sequence and a highly repetitive sequence in the subterminal region essential for transposition. Genetics 174, 639–649, doi:10.1534/genetics.106.060244 (2006).

26. Pounds, S. et al. A procedure to statistically evaluate agreement of differential expression for cross-species genomics. Bioinformatics 27, 2098–2103, doi:10.1093/bioinformatics/btr362 (2011).

27. White, R. J. et al. A high-resolution mRNA expression time course of embryonic development in zebrafish. Elife 6, doi:10.7554/eLife.30860 (2017).

28. Donaldson, J. G. Multiple roles for Arf6: sorting, structuring, and signaling at the plasma membrane. J Biol Chem 278, 41573–41576, doi:10.1074/jbc.R300026200 (2003).

29. Howe, K. et al. The zebrafish reference genome sequence and its relationship to the human genome. Nature 496, 498–503, doi:10.1038/nature12111 (2013).

30. McConnell, A. M., Noonan, H. R. & Zon, L. I. Reeling in the Zebrafish Cancer Models. Annual Review of Cancer Biology 5, 331–350, doi:10.1146/annurev-cancerbio-051320-014135 (2021).

31. Amatruda, J. F. Modeling the developmental origins of pediatric cancer to improve patient outcomes. Dis Model Mech 14, doi:10.1242/dmm.048930 (2021).

32. Mielcarek, M., Gunther, S., Kruger, M. & Braun, T. VITO-1, a novel vestigial related protein is predominantly expressed in the skeletal muscle lineage. Mech Dev 119 Suppl 1, S269–274, doi:10.1016/s0925-4773(03)00127-8 (2002).

33. Honda, M. et al. Vestigial-like 2 contributes to normal muscle fiber type distribution in mice. Sci Rep 7, 7168, doi:10.1038/s41598-017-07149-0 (2017).

34. Honda, M., Tsuchimochi, H., Hitachi, K. & Ohno, S. Transcriptional cofactor Vgll2 is required for functional adaptations of skeletal muscle induced by chronic overload. J Cell Physiol, doi:10.1002/jcp.28239 (2019).

35. Hashimoto, S. et al. Requirement for Arf6 in breast cancer invasive activities. Proc Natl Acad Sci U S A 101, 6647–6652, doi:10.1073/pnas.0401753101 (2004).

36. Stepicheva, N. A., Dumas, M., Kobi, P., Donaldson, J. G. & Song, J. L. The small GTPase Arf6 regulates sea urchin morphogenesis. Differentiation 95, 31–43, doi:10.1016/j.diff.2017.01.003 (2017).

37. Suzuki, T. et al. Crucial role of the small GTPase ARF6 in hepatic cord formation during liver development. Mol Cell Biol 26, 6149–6156, doi:10.1128/MCB.00298-06 (2006).

38. Chen, E. H., Pryce, B. A., Tzeng, J. A., Gonzalez, G. A. & Olson, E. N. Control of myoblast fusion by a guanine nucleotide exchange factor, loner, and its effector ARF6. Cell 114, 751–762, doi:10.1016/s0092-8674(03)00720-7 (2003).

39. Morishige, M. et al. GEP100 links epidermal growth factor receptor signalling to Arf6 activation to induce breast cancer invasion. Nat Cell Biol 10, 85–92, doi:10.1038/ncb1672 (2008).

40. Yoo, J. H. et al. ARF6 Is an Actionable Node that Orchestrates Oncogenic GNAQ Signaling in Uveal Melanoma. Cancer Cell 29, 889–904, doi:10.1016/j.ccell.2016.04.015 (2016).

41. Yoo, J. H. et al. The Small GTPase ARF6 Activates PI3K in Melanoma to Induce a Prometastatic State. Cancer Res 79, 2892–2908, doi:10.1158/0008-5472.CAN-18-3026 (2019).

42. Oka, S., Uramoto, H., Shimokawa, H., Yamada, S. & Tanaka, F. Epidermal growth factor receptor-GEP100-Arf6 axis affects the prognosis of lung adenocarcinoma. Oncology 86, 263–270, doi:10.1159/000360089 (2014).

43. Hashimoto, S. et al. Lysophosphatidic acid activates Arf6 to promote the mesenchymal malignancy of renal cancer. Nat Commun 7, 10656, doi:10.1038/ncomms10656 (2016).

44. Yamauchi, Y., Miura, Y. & Kanaho, Y. Machineries regulating the activity of the small GTPase Arf6 in cancer cells are potential targets for developing innovative anti-cancer drugs. Adv Biol Regul 63, 115–121, doi:10.1016/j.jbior.2016.10.004 (2017).

45. Berghmans, S. et al. tp53 mutant zebrafish develop malignant peripheral nerve sheath tumors. Proc Natl Acad Sci U S A 102, 407–412, doi:10.1073/pnas.0406252102 (2005).

46. Kendall, G. C. & Amatruda, J. F. Zebrafish as a Model for the Study of Solid Malignancies. Methods Mol Biol 1451, 121–142, doi:10.1007/978-1-4939-3771-4_9 (2016).

47. Kwan, K. M. et al. The Tol2kit: a multisite gateway-based construction kit for Tol2 transposon transgenesis constructs. Dev Dyn 236, 3088–3099, doi:10.1002/dvdy.21343 (2007).

48. Mosimann, C. et al. Ubiquitous transgene expression and Cre-based recombination driven by the ubiquitin promoter in zebrafish. Development 138, 169–177, doi:10.1242/dev.059345 (2011).

49. Berger, J. & Currie, P. D. 503unc, a small and muscle-specific zebrafish promoter. Genesis 51, 443–447, doi:10.1002/dvg.22385 (2013).

50. Kawakami, K. et al. A transposon-mediated gene trap approach identifies developmentally regulated genes in zebrafish. Dev Cell 7, 133–144, doi:10.1016/j.devcel.2004.06.005 (2004).

51. Provost, E., Rhee, J. & Leach, S. D. Viral 2A peptides allow expression of multiple proteins from a single ORF in transgenic zebrafish embryos. Genesis 45, 625–629, doi:10.1002/dvg.20338 (2007).

52. Langmead, B. & Salzberg, S. L. Fast gapped-read alignment with Bowtie 2. Nat Methods 9, 357–359, doi:10.1038/nmeth.1923 (2012).

53. Dobin, A. et al. STAR: ultrafast universal RNA-seq aligner. Bioinformatics 29, 15–21, doi:10.1093/bioinformatics/bts635 (2013).

54. Li, H. et al. The Sequence Alignment/Map format and SAMtools. Bioinformatics 25, 2078–2079, doi:10.1093/bioinformatics/btp352 (2009).

55. Anders, S., Pyl, P. T. & Huber, W. HTSeq--a Python framework to work with high-throughput sequencing data. Bioinformatics 31, 166–169, doi:10.1093/bioinformatics/btu638 (2015).

56. Gentleman, R. C. et al. Bioconductor: open software development for computational biology and bioinformatics. Genome Biol 5, R80, doi:10.1186/gb-2004-5-10-r80 (2004).

57. Robinson, M. D., McCarthy, D. J. & Smyth, G. K. edgeR: a Bioconductor package for differential expression analysis of digital gene expression data. Bioinformatics 26, 139–140, doi:10.1093/bioinformatics/btp616 (2010).

58. McCarthy, D. J., Chen, Y. & Smyth, G. K. Differential expression analysis of multifactor RNA-Seq experiments with respect to biological variation. Nucleic Acids Res 40, 4288–4297, doi:10.1093/nar/gks042 (2012).

59. Kanehisa, M., Furumichi, M., Tanabe, M., Sato, Y. & Morishima, K. KEGG: new perspectives on genomes, pathways, diseases and drugs. Nucleic Acids Res 45, D353–D361, doi:10.1093/nar/gkw1092 (2017).

60. Bray, N. L., Pimentel, H., Melsted, P. & Pachter, L. Near-optimal probabilistic RNA-seq quantification. Nat Biotechnol 34, 525–527, doi:10.1038/nbt.3519 (2016).

61. Cancer Genome Atlas Research Network. Electronic address, e. d. s. c. & Cancer Genome Atlas Research, N. Comprehensive and Integrated Genomic Characterization of Adult Soft Tissue Sarcomas. Cell 171, 950–965 e928, doi:10.1016/j.cell.2017.10.014 (2017).

62. Crompton, B. D. et al. The genomic landscape of pediatric Ewing sarcoma. Cancer Discov 4, 1326–1341, doi:10.1158/2159-8290.CD-13-1037 (2014).

63. Xu, L. et al. Integrative Bayesian Analysis Identifies Rhabdomyosarcoma Disease Genes. Cell Rep 24, 238–251, doi:10.1016/j.celrep.2018.06.006 (2018).

64. Babicki, S. et al. Heatmapper: web-enabled heat mapping for all. Nucleic Acids Res 44, W147–153, doi:10.1093/nar/gkw419 (2016).

65. Huang da, W., Sherman, B. T. & Lempicki, R. A. Systematic and integrative analysis of large gene lists using DAVID bioinformatics resources. Nat Protoc 4, 44–57, doi:10.1038/nprot.2008.211 (2009).

66. Huang da, W., Sherman, B. T. & Lempicki, R. A. Bioinformatics enrichment tools: paths toward the comprehensive functional analysis of large gene lists. Nucleic Acids Res 37, 1–13, doi:10.1093/nar/gkn923 (2009).

