## Supplemental Figures and Tables for "*VGLL2-NCOA2* leverages developmental programs for pediatric sarcomagenesis"

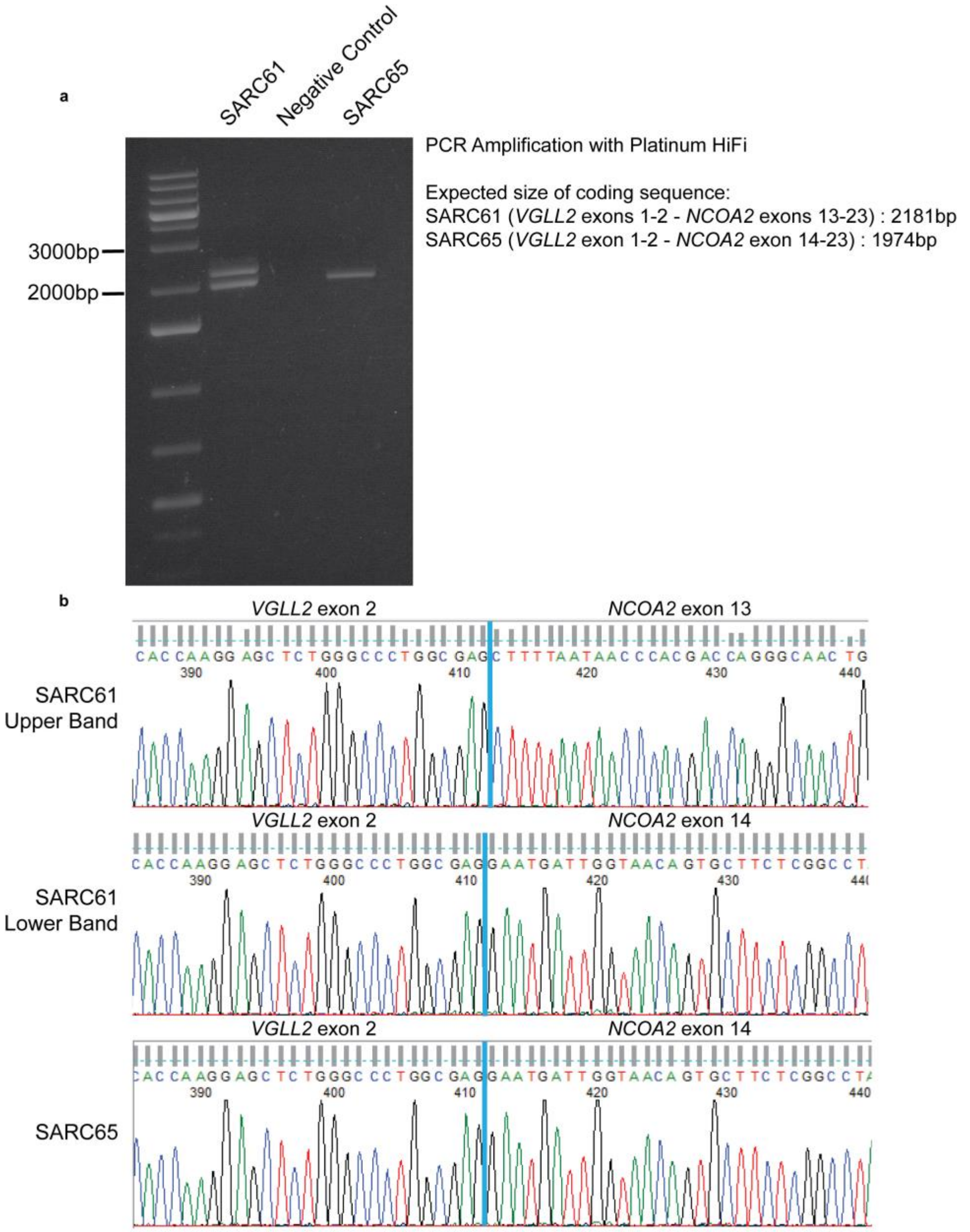


**Supplemental Figure 1. Cloning of the _VGLL2-NCOA2_ fusion from primary patient tumors.**

A) RT-PCR showing two patient samples amplified with a forward primer in *VGLL2* and a reverse primer in *NCOA2*. B) Sanger sequencing analysis of the junctions in each band of A.

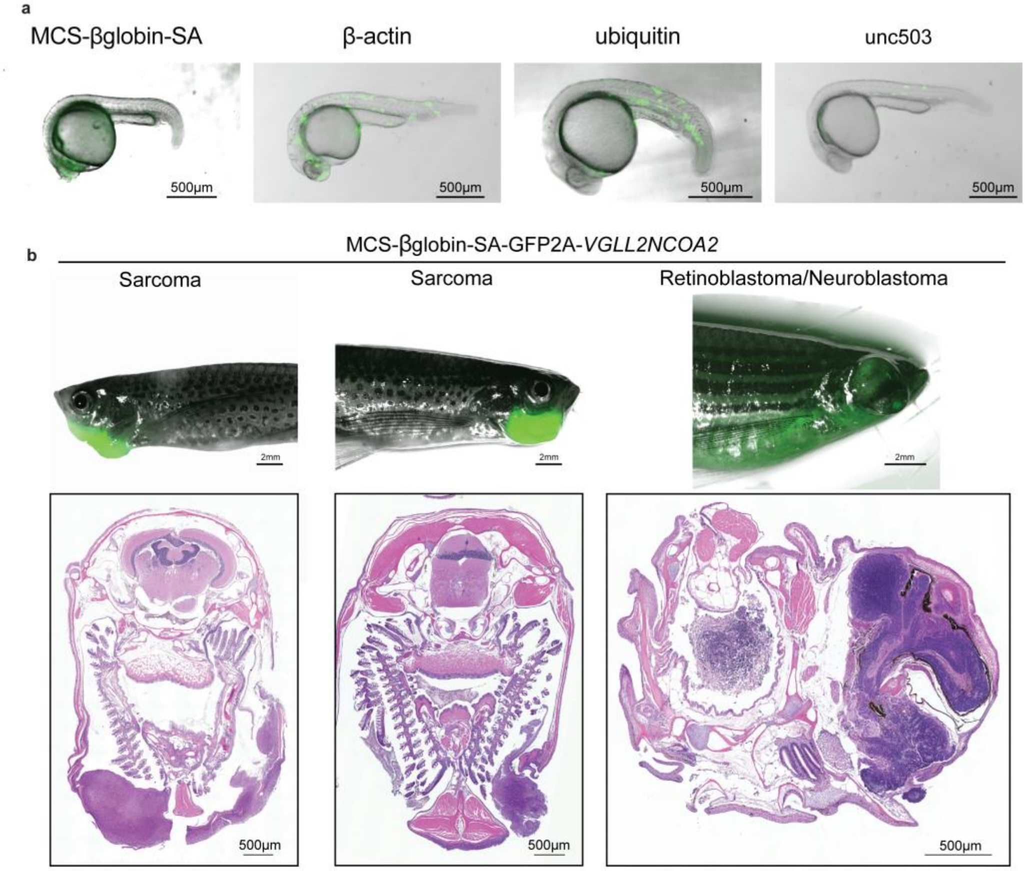


c

| Promoter | Number of GFP+ Adult Fish |
|---|---|
| MCS-βglobin-SA | 6 / 42 |
| β-actin | 3 / 50 |
| ubiquitin | 3 / 36 |
| unc503 | 1 / 51 |

**Supplemental Figure 2. Presentation of zebrafish injected with *VGLL2-NCOA2* under control of alternate promoters.**

A) Embryos injected at the single cell stage with GFP-tagged *VGLL2-NCOA2* under control of promoters indicated: MCS-beta-globin-Splice Acceptor, beta-actin, ubiquitin, and unc503. Images taken at 24 hours post fertilization. B) Images of MCS-beta-globin-SA driven *VGLL2-NCOA2* tumors, accompanied by H&E stains of transverse sections. C) Summary table of numbers of GFP positive fish and total number of fish screened for each promoter.

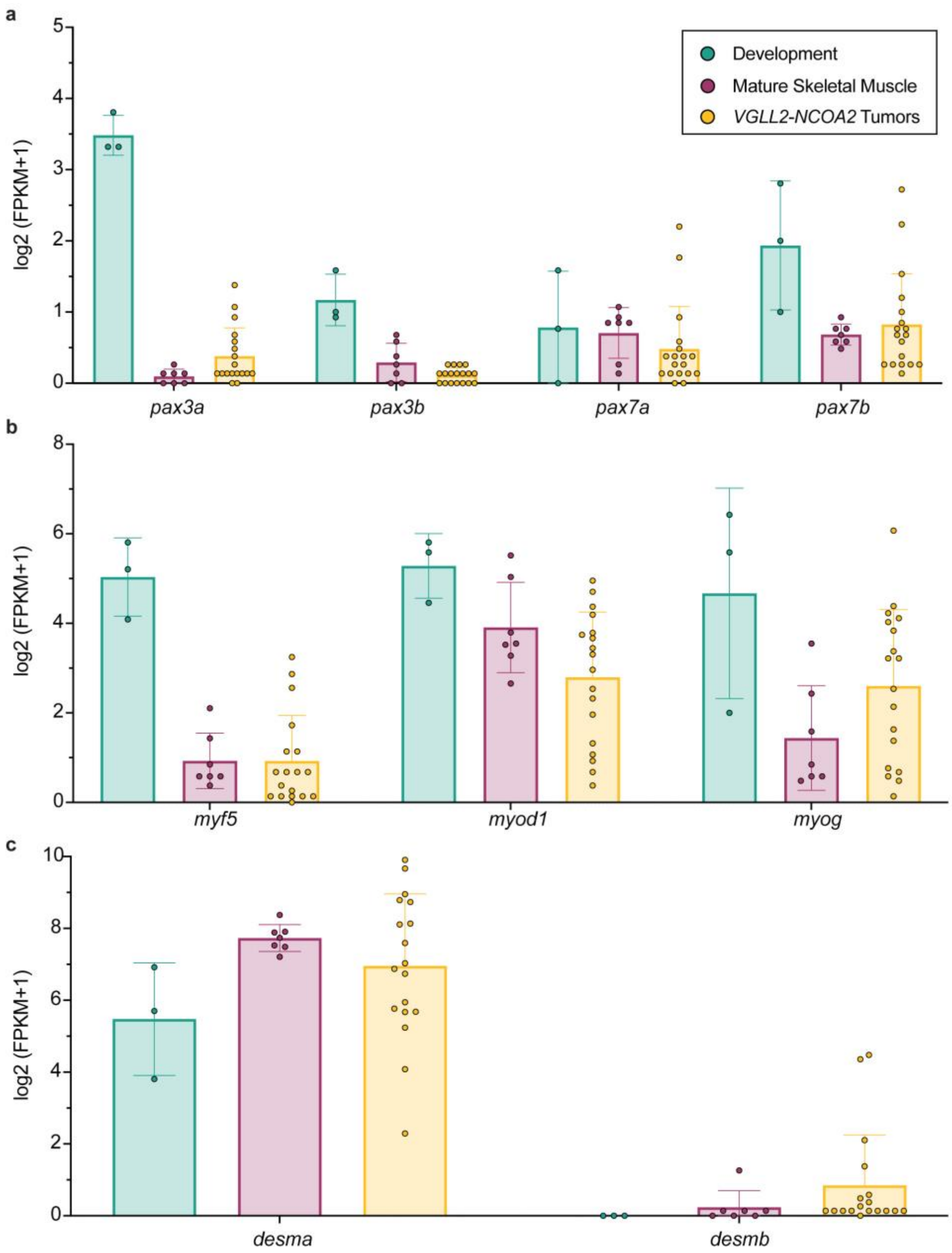


**Supplemental Figure 3. RNA expression of muscle regulatory factors in development, skeletal muscle, and *VGLL2-NCOA2* tumors.**

Data as in Figure 4: RNA sequencing was performed on n=18 zebrafish *VGLL2-NCOA2* tumors, n=7 zebrafish mature skeletal muscle samples and n=3 pooled samples of larval zebrafish from segmentation timepoints at 1-4 somites, 14-19 somites, and 20-25 somites (10.33, 16, and 19 hours post fertilization at 28°C, respectively). Each point is a sample. A) Expression of pax transcription factors. Plot of FPKM values for pax family members, pax3 and pax7 genes, both of which are duplicated in zebrafish. B) Expression of muscle regulator factors. Plot of FPKM values from muscle regulatory factors *myf5*, *myod*, and *myog*. C) Expression of rhabdomyosarcoma diagnostic marker, desmin. Plot of FPKM values for desmin (*desma* and *desmb*), which is duplicated in zebrafish. All genes are expressed in *VGLL2-NCOA2* tumors but are statistically insignificant as compared to skeletal muscle or developmental samples.

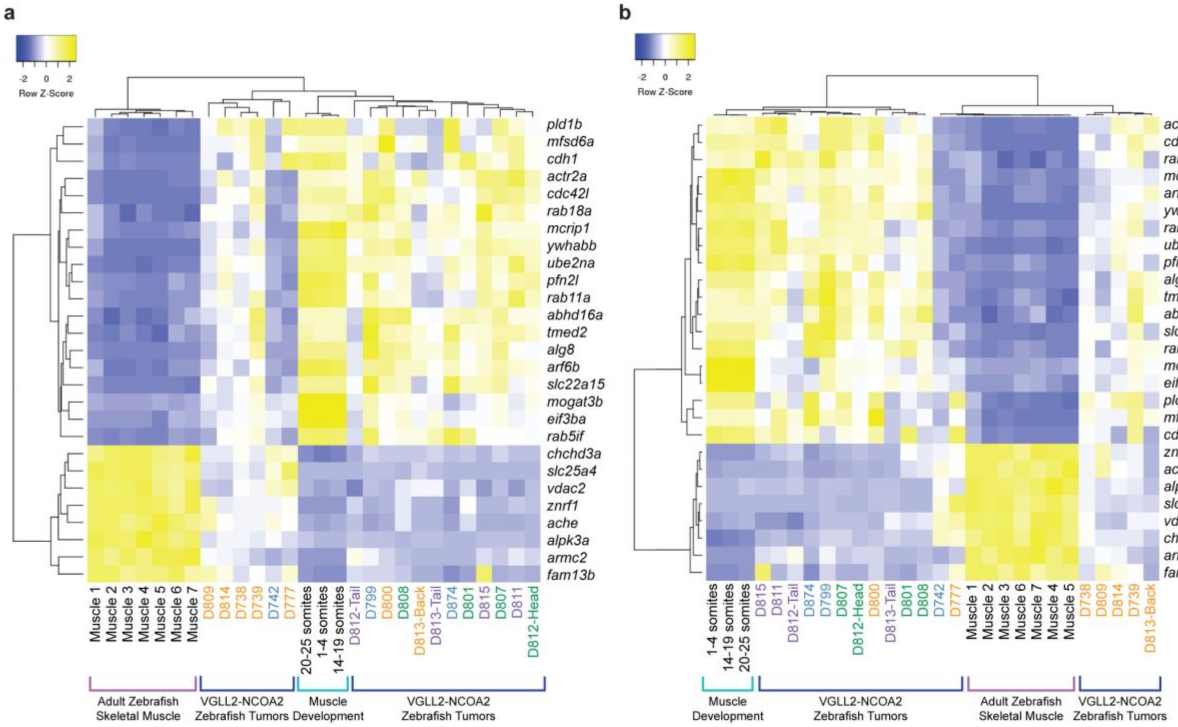


**Supplemental Figure 4. Alternate clustering analyses of *VGLL2-NCOA2* tumors, development, and skeletal muscle.**

Data as in Figure 4: Heat maps plotting the FPKM of the n=27 differentially expressed genes from Figure 4A-D. The samples are clustered by row and column using the average linkage clustering method. Tumors are colored based on location on the fish: back tumors in orange, ventral tumors in blue, head tumors in green and tail tumors in purple. A) Kendall's Tau distance measurement method was utilized. B) Pearson distance measurement method was utilized.

```
zebrafish_arf6a      MGKMLSKIFGNKEMRILMLGLDAAGKTTILYKLKLGQSVTTIPTVGFNVETVTYKNVKFN      60
zebrafish_arf6b      MGKMLSKIFGNKEMRILMLGLDAAGKTTILYKLKLGQSVTTIPTVGFNVETVTYKNVKFN      60
human_ARF6           MGKVLSKIFGNKEMRILMLGLDAAGKTTILYKLKLGQSVTTIPTVGFNVETVTYKNVKFN      60
                     ***:********************************************************

zebrafish_arf6a      VWDVGGQDKIRPLWRHYYTGTQGLIFVVDCADRDRIDEARQELHRIINDREMRDAIILIF      120
zebrafish_arf6b      VWDVGGQDKIRPLWRHYYTGTQGLIFVVDCADRDRIDEARQELHRIINDREMRDAIILIF      120
human_ARF6           VWDVGGQDKIRPLWRHYYTGTQGLIFVVDCADRDRIDEARQELHRIINDREMRDAIILIF      120
                     ************************************************************

zebrafish_arf6a      ANKQDLPDAMKPHEIQEKLGLTRIRDRNWYVQPSCATTGDGLYEGLTWLTSNYKS           175
zebrafish_arf6b      ANKQDLPDAMKPHEIQEKLGLTRIRDRNWYVQPSCATTGDGLYEGLTWLTSNYKS           175
human_ARF6           ANKQDLPDAMKPHEIQEKLGLTRIRDRNWYVQPSCATSGDGLYEGLTWLTSNYKS           175
                     *************************************:*****************
```

**Supplemental Figure 5. Human and zebrafish ARF6 orthologs are highly conserved.** Sequence alignment of human ARF6 and zebrafish arf6a and arf6b protein sequences using Clustal O(1.2.4). An asterisk indicates perfect alignment and a colon indicates strong similarity. Zebrafish arf6a and arf6b have 100% identity. Human ARF6 and zebrafish arf6a/b sequences share 99% identity (173/175 amino acids) and 100% similarity (175/175 amino acids).

| Gender | Age at Diagnosis | Primary tumor site | Metastases at diagnosis | Tissue | Histology | Immunohistochemistry | Initial diagnosis | Relapse | Identified Fusion | CGH array |
|---|---|---|---|---|---|---|---|---|---|---|
| M | 4.5 years | Chest | - | Soft tissue | | Desmin + (focal), Myogenin + (focal), INI1+, Ki67 30% | Embryonal RMS | NA | *VGLL2* ex2-*NCOA2* ex14 | |
| F | 2 months | Left arm | - | Soft tissue | | Myogenin + (focal), Ki67 30% | Immature embryonal RMS | NA | *VGLL2* ex2-*NCOA2* ex14 | Gain chr17 |
| ? | Therapeutic abortion 29 weeks | Cervical | Bone marrow, lungs, liver | Soft tissue | Mesenchymal tumor cells with rhabdomyoblastic differentiation | ? | ? | | *VGLL2* ex2-*NCOA2* ex14 | |
| M | 2 days | Left shoulder | - | Soft tissue | | CD34+ (focal) | Embryonal RMS | Metastatic relapse at 2 years (lungs, bone, meninges) | *VGLL2* ex2-*NCOA2* ex13 | |
| F | 6 days | Left arm | - | Soft tissue | | Ki67 25%, a-actin + (focal) | Congenital fibrosarcoma or embryonal RMS | Local progression at 3 months | *VGLL2* ex2-*NCOA2* ex 14 | Flat profile |

**Supplemental Table 1. Clinical presentation and diagnosis of patients with *VGLL2-NCOA2* tumors from Watson et al 2018 [4].**

| Tumor Location | Number of Fish | Average Age of Sac | Median Age of Sac | Range of Age at Sac |
|---|---|---|---|---|
| Tail | 33 | 67.58 | 55 | 30-358 |
| Back | 17 | 106.00 | 53 | 40-393 |
| Head/Eye | 15 | 64.14 | 51.5 | 30-112 |
| Ventral | 7 | 138.14 | 119 | 50-262 |
| Head and Tail | 2 | 47.5 | 47.5 | 47-48 |
| Back and Tail | 1 | 47 | 47 | 47 |
| Total | 75 | 81.61 | 55 | 30-393 |

**Supplemental Table 2. Tumor onset data for CMV driven *VGLL2-NCOA2* zebrafish tumors.** Displayed are the tumor locations, number of fish for each location, and the average, median, and range of the age at sacrifice (sac) in days. The age at sacrifice is defined as the day fish were euthanized and samples collected due to tumor burden and failure to thrive.

| Gene Target | Organism | Primer Name and #s | Forward Primer | Reverse Primer |
|---|---|---|---|---|
| *VGLL2-NCOA2* | Human | 5659-5660 | GCTGCGTCCTCTTCACTTATT | GCATAGTAGGCCGAGAAGCA |
| *VGLL2-NCOA2* | Cloning Human | VGLL2_BamH1_FWD / NCOA2_Xho1_REV | ATCGGGATCCATGAGCTGCTGGATGTT | ATCGCTCGAGTCAGCAATATTTCCGTGTTG |
| *myod1* | Zebrafish | 6278-6279 | ATCCAACTGCTCTGATGGC | AGACAATCCAAACTCGACACC |
| *myog* | Zebrafish | 6280-6281 | GCTATACAGTACATCGAGAGGC | AGAGCCCTGATCACTAGAGG |
| *desma* | Zebrafish | 6361-6362 | GCAGTGAACAAGAATAACGAGG | CACTCATTTGCCTCCTCAGAG |
| *arf6a* | Zebrafish | 6888-6889 | ACGCCATTATCCTCATCTTCG | CCAGCTTCTCCTGTATCTCATG |
| *arf6b* | Zebrafish | 6890-6891 | TCAAGTTCAACGTGTGGGAC | GTCTATGCGATCTCTATCTGCG |
| *rab11a* | Zebrafish | 6880-6881 | GCCTCCACTTTCCTTCACATTC | GTTACTCTTCCCCACACCAG |
| *rab18a* | Zebrafish | 6882-6883 | ATTGGCGAAAGTGGAGTAGG | TTGCTCGGTTTCCATCTACAG |
| *ywhabb* | Zebrafish | 6874-6875 | ATATTTATCTGAGGTAGCATCCGG | GGCAAGACCCAACCGTATAG |
| *cdh1* | Zebrafish | 6870-6871 | AGGAGAAGTTTATTTCCAGACCTG | AGCCTGTTATATTGAGCCAGTC |
| *gapdh* | Zebrafish | 5715-5716 | GTGGCCATCAATGACCCATTC | CAATGACCAGTTTGCCGCCTTC |
| *rpl13a* | Zebrafish | 3809-3810 | CGGTCGTCTTTCCGCTATT | TTCCAGAGATGTTGATACCCTCAC |

**Supplemental Table 3. List of oligonucleotide primer sequences.**

| Sample ID Curie | Sample ID | Tissue Type | Zebrafish Genetic Background | Age at Sac (Days) | Transgene | Location Sequenced |
|---|---|---|---|---|---|---|
| A667T19 | D738 | VGLL2-NCOA2 Tumor | tp53M214K mutant | 44 | ptz876 cmv-GFP2A-VGLL2NCOA2 injected | Institut Curie |
| A667T20 | D739 | VGLL2-NCOA2 Tumor | Wildtype (AB/TL) | 49 | ptz876 cmv-GFP2A-VGLL2NCOA2 injected | Institut Curie |
| A667T21 | D742 | VGLL2-NCOA2 Tumor | tp53M214K mutant | 51 | ptz876 cmv-GFP2A-VGLL2NCOA2 injected | Institut Curie |
| A667T22 | D777 | VGLL2-NCOA2 Tumor | Wildtype (AB/TL) | 50 | ptz876 cmv-GFP2A-VGLL2NCOA2 injected | Institut Curie |
| A667T23 | D799 | VGLL2-NCOA2 Tumor | Wildtype (AB/TL) | 75 | ptz876 cmv-GFP2A-VGLL2NCOA2 injected | Institut Curie |
| A667T24 | D800 | VGLL2-NCOA2 Tumor | Wildtype (AB/TL) | 75 | ptz876 cmv-GFP2A-VGLL2NCOA2 injected | Institut Curie |
| A667T25 | D801 | VGLL2-NCOA2 Tumor | Wildtype (AB/TL) | 75 | ptz876 cmv-GFP2A-VGLL2NCOA2 injected | Institut Curie |
| A667T26 | D807 | VGLL2-NCOA2 Tumor | Wildtype (AB/TL) | 100 | ptz876 cmv-GFP2A-VGLL2NCOA2 injected | Institut Curie |
| A667T27 | D808 | VGLL2-NCOA2 Tumor | Wildtype (AB/TL) | 44 | ptz876 cmv-GFP2A-VGLL2NCOA2 injected | Institut Curie |
| A667T28 | D813-Head | VGLL2-NCOA2 Tumor | Wildtype (AB/TL) | 47 | ptz876 cmv-GFP2A-VGLL2NCOA2 injected | Institut Curie |
| A667T29 | D813-Tail | VGLL2-NCOA2 Tumor | Wildtype (AB/TL) | 47 | ptz876 cmv-GFP2A-VGLL2NCOA2 injected | Institut Curie |
| none | D809 | VGLL2-NCOA2 Tumor | Wildtype (AB/TL) | 44 | ptz876 cmv-GFP2A-VGLL2NCOA2 injected | DNALinks |
| none | D811 | VGLL2-NCOA2 Tumor | Wildtype (AB/TL) | 47 | ptz876 cmv-GFP2A-VGLL2NCOA2 injected | DNALinks |
| none | D812-head | VGLL2-NCOA2 Tumor | Wildtype (AB/TL) | 47 | ptz876 cmv-GFP2A-VGLL2NCOA2 injected | DNALinks |
| none | D812-tail | VGLL2-NCOA2 Tumor | Wildtype (AB/TL) | 47 | ptz876 cmv-GFP2A-VGLL2NCOA2 injected | DNALinks |
| none | D814 | VGLL2-NCOA2 Tumor | Wildtype (AB/TL) | 47 | ptz876 cmv-GFP2A-VGLL2NCOA2 injected | DNALinks |
| none | D815 | VGLL2-NCOA2 Tumor | Wildtype (AB/TL) | 47 | ptz876 cmv-GFP2A-VGLL2NCOA2 injected | DNALinks |
| none | D874 | VGLL2-NCOA2 Tumor | Wildtype (AB/TL) | 145 | ptz876 cmv-GFP2A-VGLL2NCOA2 injected | DNALinks |
| A374-A375T23 | Muscle2 | Normal Back Skeletal Muscle | Wildtype (AB/TL) | 258 | - | Institut Curie |
| A374-A375T24 | Muscle3 | Normal Back Skeletal Muscle | Wildtype (AB/TL) | 258 | - | Institut Curie |
| none | D948 muscle | Normal Back Skeletal Muscle | Wildtype (AB/TL) | 118 | - | DNALinks |
| none | D949 muscle | Normal Back Skeletal Muscle | Wildtype (AB/TL) | 118 | - | DNALinks |
| none | D950 muscle | Normal Back Skeletal Muscle | Wildtype (AB/TL) | 118 | - | DNALinks |
| none | D951 muscle | Normal Back Skeletal Muscle | Wildtype (AB/TL) | 118 | - | DNALinks |
| none | D952 muscle | Normal Back Skeletal Muscle | Wildtype (AB/TL) | 118 | - | DNALinks |

**Supplemental Table 4. Zebrafish *VGLL2-NCOA2* tumor and mature skeletal muscle sample details used for RNA-seq.**
